# Dispersal alters the nature and scope of sexually antagonistic variation

**DOI:** 10.1101/2020.05.05.080010

**Authors:** Ewan O Flintham, Vincent Savolainen, Charles Mullon

## Abstract

Intralocus sexual conflict, or sexual antagonism, occurs when alleles have opposing fitness effects in the two sexes. Previous theory suggests that sexual antagonism is a driver of genetic variation by generating balancing selection. However, most of these studies assume that populations are well-mixed, neglecting the effects of spatial subdivision. Here we use mathematical modelling to show that limited dispersal changes evolution at sexually antagonistic autosomal and X-linked loci due to inbreeding and sex-specific kin competition. We find that if the sexes disperse at different rates, kin competition within the philopatric sex biases intralocus conflict in favour of the more dispersive sex. Furthermore, kin competition diminishes the strength of balancing selection relative to genetic drift, reducing genetic variation in small subdivided populations. Meanwhile, by decreasing heterozygosity, inbreeding reduces the scope for sexually antagonistic polymorphism due to nonadditive allelic effects, and this occurs to a greater extent on the X-chromosome than autosomes. Overall, our results indicate that spatial structure is a relevant factor in predicting where sexually antagonistic alleles might be observed. We suggest that sex-specific dispersal ecology and demography can contribute to interspecific and intragenomic variation in sexual antagonism.

## 1 Introduction

Due to the different reproductive roles of males and females, many traits are selected in different directions in the two sexes (Parker et al., 1979). Responding to these divergent selection pressures, however, is not straightforward. Because the sexes share a large part of their genomes and traits are typically determined by the same genes, homologous traits in males and females tend to be genetically correlated (Poissant et al., 2010). Opposing selection pressures on the two sexes therefore lead to a genetic tug-of-war, whereby some alleles are favoured in one sex but disfavoured in the other (Connallon and Clark, 2014). This tug-of-war, also known as intralocus sexual conflict or sexual antagonism (Chippindale et al., 2001; Brommer et al., 2007; Mainguy et al., 2009; Foerster et al., 2007; Svensson et al., 2009; Bonduriansky and Chenoweth, 2009; Lewis et al., 2011), can result in balancing selection and the long-term maintenance of polymorphism, within and even between species (e.g. Ruzicka et al., 2019; Eyer et al., 2019).

Mathematical population genetics models have helped elucidate the conditions that favour the emergence and maintenance of sexually antagonistic variation (e.g. Owen, 1953; Kidwell et al., 1977; Rice, 1984; Albert and Otto, 2005; Gavrilets and Rice, 2006; Fry, 2010; Arnqvist, 2011; Connallon and Clark, 2012; Jordan and Charlesworth, 2012; Mullon et al., 2012; Jaquiéry et al., 2013; Harts et al., 2014; Jordan and Connallon, 2014; Tazzyman and Abbott, 2015; de Vries and Caswell, 2019; Kasimatis et al., 2019; Connallon et al., 2019; Hitchcock and Gardner, 2020; Ruzicka and Connallon, 2020). Firstly, models have highlighted the role of dominance, i.e. non-additive effects of alleles, in facilitating balancing selection at sexually antagonistic loci on both autosomes and sex-chromosomes (Kidwell et al., 1977; Rice, 1984; Fry, 2010; Spencer and Priest, 2016). In line with this prediction for autosomes, a number of empirical studies have found that loci underlying intralocus sexual conflict often exhibit sex differences in dominance that increase the fitness of both male and female heterozygotes (referred to as “dominance reversal”, Hager et al., 2008; Barson et al., 2015; Grieshop and Arnqvist, 2018; Pearse et al., 2019). Secondly, stochastic models have emphasised the importance of genetic drift in countering the effects of sexually antagonistic selection in finite populations. In fact, because sexually antagonistic selection is relatively weak, its effects on polymorphism can be easily negated by genetic drift (Connallon and Clark, 2012; Mullon et al., 2012).

A common assumption to the vast majority of models investigating sexually antagonistic variation is that populations are well-mixed or that individuals compete at random (Owen, 1953; Kidwell et al., 1977; Rice, 1984; Albert and Otto, 2005; Gavrilets and Rice, 2006; Fry, 2010; Connallon and Clark, 2012; Jordan and Charlesworth, 2012; Mullon et al., 2012; Jaquiéry et al., 2013; Jordan and Connallon, 2014; de Vries and Caswell, 2019; Ruzicka and Connallon, 2020). In reality, most natural populations are spatially structured or subdivided, simply due to the physical constraints of movement that lead to limited dispersal (Clobert et al., 2001). Where dispersal is limited, individuals interacting with one another are more likely to share common alleles than individuals sampled at random in the population (Rousset, 2004). Studies of sexual antagonism in a spatial context have so far ignored the effects of such genetic correlations (e.g., by assuming patches of infinite size, Connallon et al., 2019, although see Van Cleve et al., 2010 for general population genetics model of a dioecious subdivided population applied to imprinting). Genetic structure, however, has important evolutionary implications as it leads to inbreeding and kin selection (Hamilton, 1964; Frank, 1998; Rousset, 2004; Charlesworth and Charlesworth, 2010). These effects may be particularly relevant for the fate of sexually antagonistic alleles as inbreeding influences the abundance of heterozygotes and hence how dominance affects selection at sexually antagonistic loci (Arnqvist, 2011; Jordan and Connallon, 2014; Tazzyman and Abbott, 2015; Kasimatis et al., 2019). In addition, limited dispersal influences the effective size of a population and thus the strength of genetic drift (Caballero, 1994; Wang and Caballero, 1999; Rousset, 2004). In spite of its potential importance, the consequences of population subdivision for sexually antagonistic variation remain poorly understood.

Here, we investigate the effects of limited dispersal on the segregation of sexually antagonistic alleles. To do this, we extend panmictic population genetics models of intralocus sexual conflict (e.g. Owen, 1953; Kidwell et al., 1977; Fry, 2010) to a subdivided population consisting of patches, or groups, interconnected by dispersal (e.g. Van Cleve et al., 2010). This model allows for sex-specific dispersal and demography (i.e. different dispersal rates for males and females, and different numbers of breeding males and females within groups) as reported in many taxa (see Trochet et al., 2016; Li and Kokko, 2019, for reviews). We examine how the interplay between selection, sex-specific ecology and genetic drift influences the segregation of sexually antagonistic alleles at autosomal and sex-linked positions.

## 2 Model

Following Van Cleve et al., 2010, we consider a dioecious population that is subdivided among patches, each composed of *n*_m_ male and *n*_f_ female adults, with the following life-cycle. (i) Within each patch, adults mate at random, producing juveniles of each sex in equal proportion. After reproduction, adults die. (ii) Each juvenile either remains in its natal patch or disperses to another randomly chosen patch (so dispersal is uniform among patches as in Wright, 1931’s island model). Juveniles disperse independently from one another, with a sex specific probability, *m*_m_ and *m*_f_ in males and females respectively. (iii) Finally, within each patch and each sex, juveniles compete to fill *n*_m_ male and *n*_f_ female breeding positions in the local mating pool and become the adults of the next generation. With dispersal occurring prior to competition, our model considers hard selection (so that patches can vary in the number of adult offspring they contribute to the next generation, Roze and Rousset, 2003; Débarre and Gandon, 2011; see Appendix A.2.6 for an analysis of our model under soft selection).

In line with previous studies of sexual antagonism (e.g. Owen, 1953; Kidwell et al., 1977; Fry, 2010), we consider a genetic locus, either on an autosome or on a sex-chromosome (X in XY species or Z in ZW species), with two segregating alleles, *a* and *A*. These alleles have opposing effects on male and female competitiveness (step (iii) of the life-cycle): allele *a* increases competitiveness when present in males but decreases competitiveness when present in females; conversely, allele *A* improves female competitiveness but comes at a cost when in males (where the cost to male and female homozygotes carrying detrimental alleles is given by *c*_m_ and *c_f_* respectively, Figure 1A). So the fittest males are homozygous *aa* while the fittest females are *AA*. In *Aa* heterozygotes, the costs of the antagonistic alleles are mediated by sex-specific dominance coefficients, *h_m_* in males and *h_f_* in females, allowing for a variety of dominance effects including dominance reversal (Figure 1B-D).

**Figure 1:**
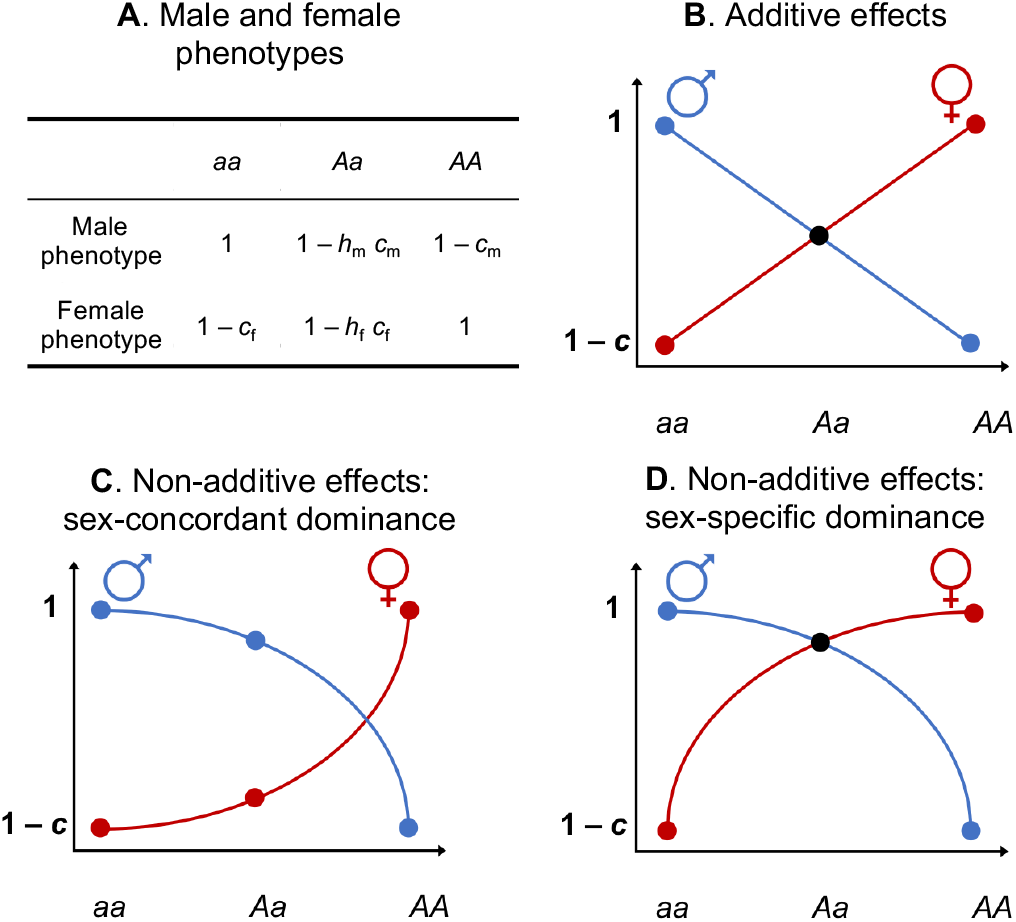
Competitiveness in each sex under different dominance scenarios. **A** shows male and female phenotypes (competitiveness) according to their genotype. **B**-**D** show how male and female competitiveness varies across genotypes depending on allele dominance effects when costs to homozygotes are equal across the sexes (*c* = *c*_m_ = *c*_f_). **B** shows additive allele effects in both sexes (co-dominance, *h*_m_ = *h*_f_ = 0.5). **C** shows non-additive effects (incomplete dominance) where the dominant and recessive alleles are the same in both sexes (*h*_m_ = 1 − *h*_f_ = 0.5). **D** shows a specific case of non-additivity and sex-specific dominance effects (*h*_m_ = 1 − *h*_f_), known as dominance reversal, whereby the detrimental allele in each sex is recessive (*h*_m_, *h*_f_ < 0.5).

## 3 Results

### 3.1 The effects of limited dispersal on sexually antagonistic selection

We first explore the effect of limited dispersal on sexually antagonistic selection by considering mathematically the change in average frequency (weighted by reproductive value, e.g. eq. 35 of Roze and Rousset, 2003, eq. 2 of Van Cleve et al., 2010) of the ‘male-detrimental’ or ‘female-beneficial’ allele *A*. For tractability, our analysis initially assumes weak selection (i.e. 0 < *c*_m_, *c*_f_ ≪ 1), large female fecundity (large enough to ignore demographic stochasticity) and an effectively infinite number of patches (see Appendix A for derivation).

#### 3.1.1 Limited dispersal introduces kin selection and inbreeding

We find that the change *Δp* (*p*) in average frequency *p* in the population of the *A* allele at an autosomal locus over one generation is

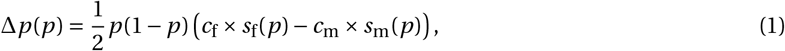

where *p* (1 − *p*) is the genetic variance in the population, *c*_f_ × *s*_f_(*p*) captures (positive) selection on *A* due to its effect in females, and -*c*_m_ × *s*_m_(*p*) captures (negative) selection on *A* due to its effect in males (see Appendices A.2.1-A.2.5 for derivation). Selection on females and males, *s*_f_(*p*) and *s*_m_(*p*), both depend on allelic frequency *p* and can be decomposed as the sum of three relevant effects,

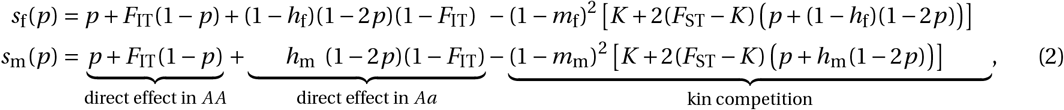

where *F*_IT_, *F*_ST_ and *K* denote probabilities that different types of genes are identical-by-descent (IBD) under neutrality (when *c*_f_ = *c*_m_ = 0), capturing various genetic consequences of limited dispersal. *F*_IT_ is the probability that the two homologous genes at an autosomal locus of one individual are IBD (which corresponds to the absolute coefficient of inbreeding in the infinite island model, Wright, 1922; Caballero, 1994; Rousset, 2004). *F*_ST_, meanwhile, is the probability that two autosomal genes in two different juveniles of the same sex and group before dispersal are IBD. This measures the degree of genetic differentiation among groups and is proportional to standard relatedness coefficients (Rousset, 2004). Due to random mating within groups, these coefficients are equal in male and female juveniles prior to dispersal. After sex-specific dispersal however, male and female genetic differentiation among groups diverge, becoming (1 − *m*_m_)^2^*F*_ST_ and (1 − *m*_f_)^2^*F*_ST_, respectively (where recall *m*_m_ and *m*_f_ are sex-specific probabilities of dispersal). Finally, *K* is the probability that the two homologous genes of a juvenile plus a third gene sampled in another juvenile of the same sex and from the same group are all IBD (also prior to dispersal). As such, (1 − *m_u_*)^2^*K* is the probability that among two competing individuals of sex *u* after dispersal, one is homozygote for a given allele and the other carries at least one IBD copy of that allele.

To understand eq. (2) and how limited dispersal influences selection, note first that in a well-mixed and randomly mating population (where *F*_IT_ = *F*_ST_ = *K* = 0), selection reduces to: *S*_f_(*p*) = *p* + (1 − *h*_f_)(1 − 2*p*) and *s*_m_(*p*) = *p* + *h*_m_(1 − 2*p*) (as found previously, e.g. eq. 2 of Mullon et al., 2012). In these baseline expressions, the term *p* captures selection on *A* due to the effects of the allele on the fitness of its bearer in homozygotes, while (1 − *h*_f_)(1 − 2*p*) and *h*_m_(1 − 2*p*) captures selection on *A* through female and male heterozygotes, respectively. Under limited dispersal, the direct effects of *A* in homozygotes (labelled “direct effect in *AA*” in eq. 2) increase to *p* + *F*_IT_(1 − *p*) in both sexes. Conversely, the direct effects of *A* in heterozygotes (labelled “direct effect in *Aa*” in eq. 2) decrease to (1 − *h*_f_)(1 − 2*p*)(1 − *F*_IT_) in females, and to *h*_m_(1 − 2*p*)(1 − *F*_IT_) in males. Selection through homozygotes is therefore more important under limited dispersal. This is because mating within groups leads to inbreeding and therefore a relative excess of homozygotes and a deficit of heterozygotes (according to the inbreeding coefficient *F*_IT_).

The remaining terms of eq. (2) (labelled “kin competition”) capture a second effect of limited dispersal: that competing individuals are more likely to carry identical gene copies than randomly sampled individuals. Such kin competition effects increase in both sexes with the probabilities that different competing individuals carry IBD genes ((1 − *m*_f_)^2^*F*_ST_ and (1 − *m*_f_)^2^*K* in females, and (1 − *m*_m_)^2^*F*_ST_ and (1 − *m*_m_)^2^*K* in males). As shown by the negative sign in front of these terms in eq. (2), kin competition effects oppose direct selection effects and therefore weaken the selective force favouring the most adaptive allele in each sex. This is because kin competition results in an individual’s reproductive success coming at the expense of genetic relatives, thus reducing the strength of selection on competitiveness.

#### 3.1.2 Additive effects: selection favours the most dispersive sex

Eq. (2) shows that where dispersal differs between males and females (*m*_f_ ≠ *m*_m_), the strength of kin competition differs among the sexes. Sex-specific dispersal therefore biases the intersexual tug-of-war over allele frequency when competition occurs after dispersal (i.e., under hard selection, see eq. A-35 in Appendix A.2.6 for soft selection). Specifically, the sex that is the most dispersal limited, and therefore experiences the greatest level of kin competition, is under weaker selection for competitiveness. This can be seen more clearly if we consider an additive locus with equally antagonistic allele effects (*h*_f_ = *h*_m_ = 1/2 and *C*_f_ = *c*_m_ = *c*). In this case, eqs. (1)-(2) reduce to

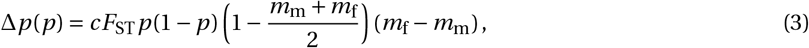

whose sign is determined by the sign of *m*_f_ − *m*_m_ (for 0 < *p* < 1;by contrast *Δp* (*p*) = 0 for all *p* for an additive locus with equally weak antagonistic effects in well-mixed populations, Fry, 2010). This means that according to the sign of *m*_f_ − *m*_m_, allele *A* or allele *a* fixes. Female beneficial *A* fixes when *m*_f_ > *m*_m_, and male beneficial *a* fixes when *m*_f_ < *m*_m_. This yields a simple rule for the fate of an additive sexually antagonistic allele: the allele that benefits the most dispersive sex will fix, and this is because the sex that disperses the most suffers less intense kin competition than the other.

#### 3.1.3 Non-additive effects: subdivision impedes the maintenance of sexually antagonistic polymorphism

In a well-mixed population, selection can favour the maintenance of sexually antagonistic polymorphism at an autosomal locus when sex-specific dominance is such that the effect of the detrimental allele in each sex is recessive, i.e. under dominance reversal (mathematically, reciprocal invasion, Δ*p*′(0) > 0 and Δ*p*′(1) < 0 where′ refers to differentiation with respect to *p*, is favoured when *h*_f_ < 1/2 and *h*_m_ < 1/2, Figure 1D, Kidwell et al., 1977; Fry, 2010 eq. 3). To investigate the effects of population subdivision on sexually antagonistic polymorphisms with sex-specific dominance, we first calculated the relevant coalescence probabilities (*F*_IT_, *F*_ST_, *K*) in eqs. (1)(2) in terms of demographic parameters following standard identity-by-descent arguments (e.g. Wang, 1997; Ramachandran et al., 2008; Van Cleve et al., 2010, for dioecious subdivided populations, our Appendix A.2.7 for derivation). We find that these coalescence probabilities are complicated expressions that depend on sexspecific dispersal (*m*_f_ and *m*_m_), as well as the number of reproducing adults in each group (*n*_f_ and *n*_m_). When dispersal among groups is low and group size is large, they simplify to

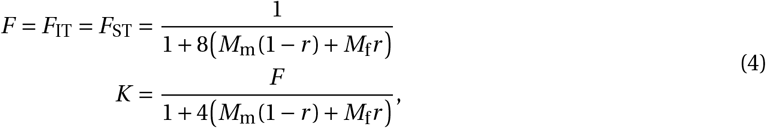

where *M*_f_ = *n*_f_*m*_f_ and *M*_m_ = *n*_m_*m*_m_ are the expected numbers of female and male immigrants in a patch at each generation, and *r* = *n*_m_/(*n*_m_ + *n*_f_) is the proportion of males among adults in a patch (i.e. the adult sex ratio;note that our expression for *F* in eq. 4 is equivalent to eq. 8 in Ramachandran et al., 2008). Unsurprisingly, eq. (4) shows that coalescence becomes more likely when there are fewer immigrants (*M*_m_ and *M*_f_ are small). But eq. (4) further tells that coalescence probabilities increase when the adult and immigrant sex ratios are both biased in a similar way (i.e. when *M*_m_ > *M*_f_ and *r* > 0.5 or when *M*_m_ < *M*_f_ and *r* < 0.5, see also Figure 2A for exact *F*_ST_ where patches are small). This is because such local bottleneck effects on the non-dispersing sex increases the coalescence of gene lineages through this sex disproportionately.

**Figure 2:**
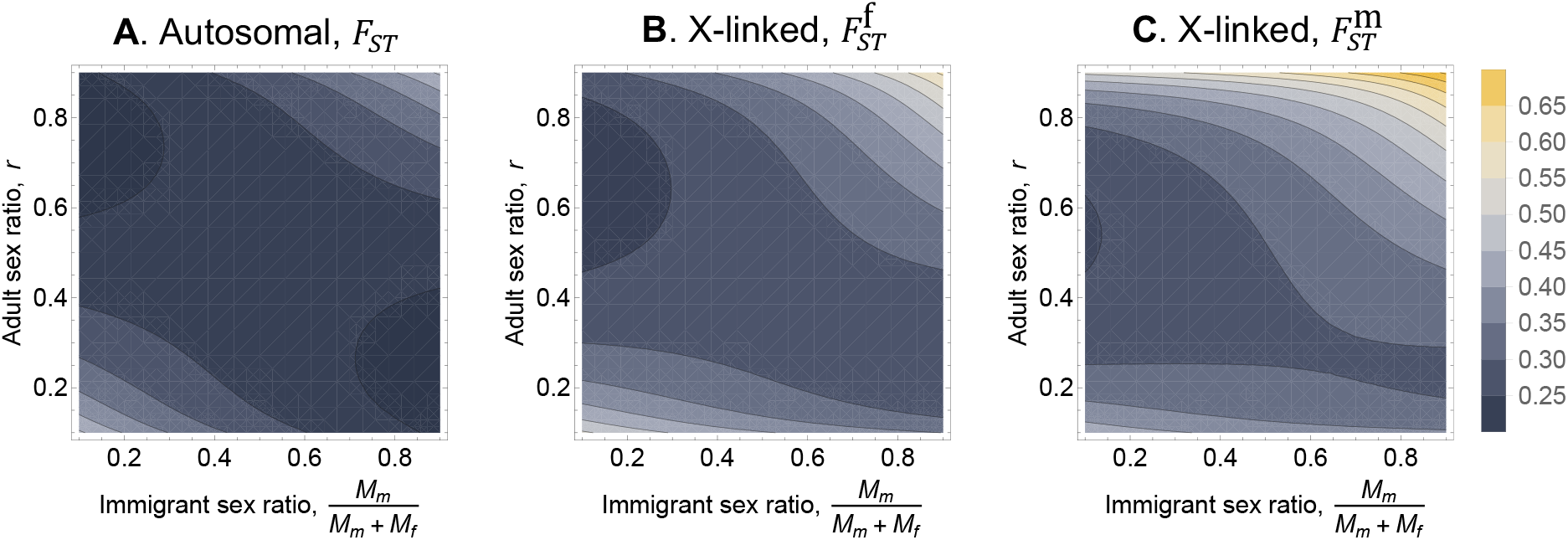
Coalescence probabilities according to adult and immigrant sex ratio. Contour plots of the probability that two neutral genes carried by two randomly sampled group neighbours are IBD when these genes are: **A.** residing on autosomes; **B.** residing on the X-chromosomes of females; **C.** residing on theX-chromosomes of males (see Appendix A.2.7 and A.3.2 for calculations;other parameters: number of immigrants per generation, *M*_m_ + *M*_f_ = 1;group size, *n*_m_ + *n*_f_ = 10, although note that group size has little incidence on *F*-statistics compared to the average numbers *M*_m_ and *M*_f_ of immigrants per generation, see eqs. 4 and 7, also Rousset, 2004).

Substituting eq. (4) (which assumes that patches are large) into eqs. (1)-(2), we find that selection favours the maintenance of sexually antagonistic polymorphism (i.e. Δ*p*’(0) > 0 and Δ*p*’(1) < 0) when

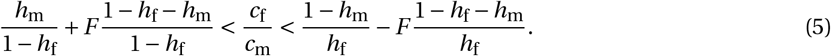

Where *F* = 0, condition eq. (5) reduces to the weak selection condition for maintenance of polymorphism in a well-mixed population (e.g. see Figure 3A; or eq. 3 in Fry, 2010 under weak selection, i.e. ignoring higher order selection effects, which using his notation are terms 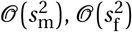 and higher). But as dispersal becomes limited in both sexes (and *F* increases), eq. (5) reveals that the conditions for polymorphism become more stringent (as the lower and upper bounds for *C*_f_/ *c*_m_ respectively increase and decrease with *F*). To check this holds more generally (i.e., for small patches) we analysed eqs.(1)-(2) numerically using exact coalescence probabilities (see Appendix A.2.7). Similarly, we find that the parameter conditions favouring polymorphism become increasingly restrictive as dispersal becomes more limited (see grey region in Figure 3A top), indicating that our conclusions also hold when patches are small (more generally, as highlighted by eqs.(1)-(2), patch size in itself has no direct effect on selection, instead what matters are genetic associations within and between individuals as summarised by *F*-statistics, see also Supplementary Figure 1). Together, these analyses show that limited dispersal decreases the scope for the maintenance of sexually antagonistic variation. This is because limited dispersal leads to inbreeding so that selection through heterozygotes, which favours sexually antagonistic polymorphism under dominance reversal, is less relevant than in well-mixed populations.

**Figure 3:**
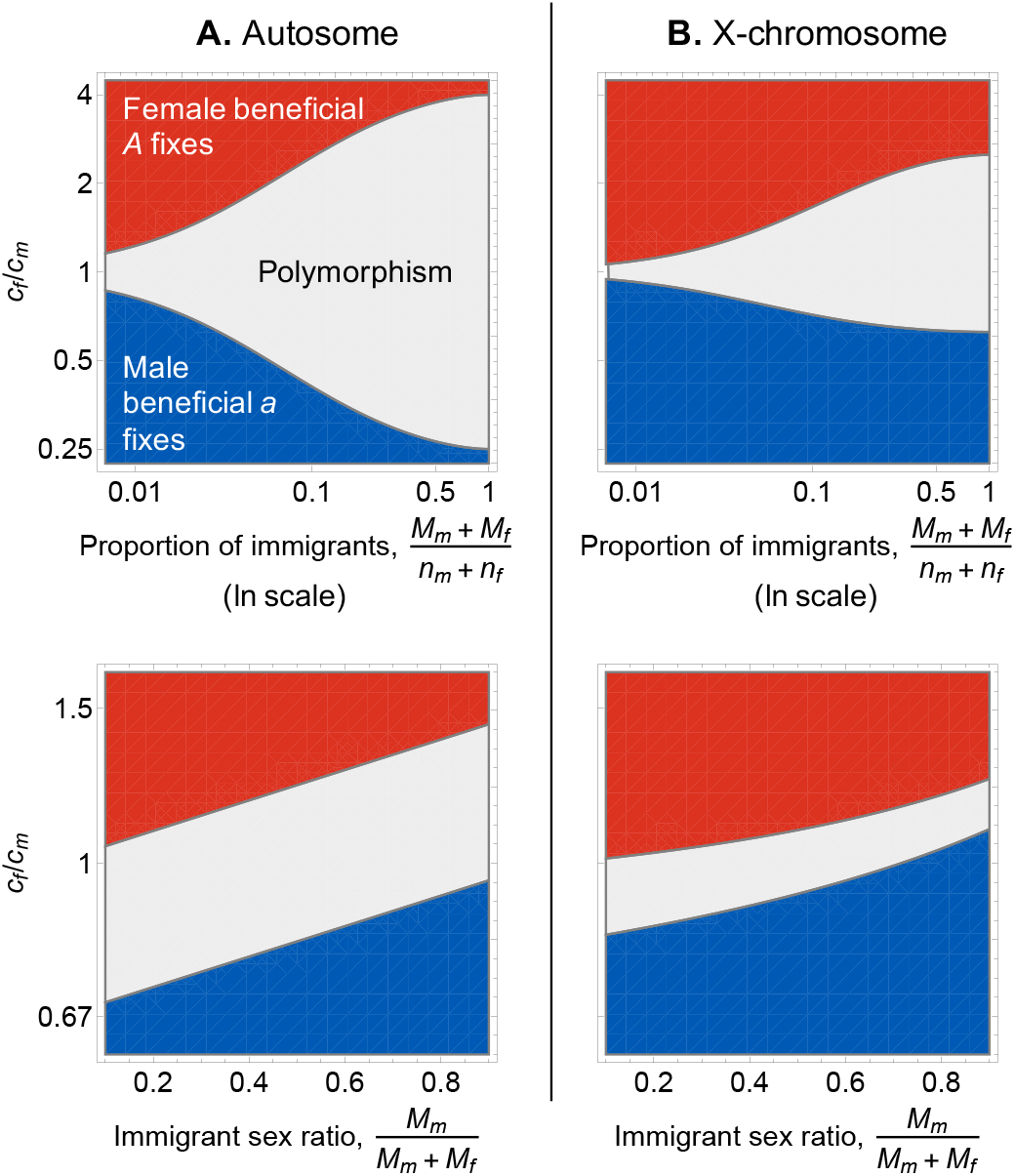
Parameters that favour the maintenance of sexually antagonistic variation. Combinations of parameters that lead to either balancing selection (grey, when Δ*p*’(0) > 0 and Δ*p’*(1) < 0), positive selection for *A* (red, when Δ*p*’(*p*) > 0 for all *p*), and positive selection for *a* (blue, when Δ*p*’(*p*) < 0 for all *p*) at **A.** an autosomal locus (computed from eqs. 1-2 with exact coalescence probabilities, see Appendix A.2.7 for calculations of these probabilities) and **B.** X-linked locus (computed from eqs. A-55-A-57 with exact coalescence probabilities, see Appendix A.3.2 for calculations). Top: selection according to ratio of homozygotic effects in females and males (*c*_m_/*c*_f_) and the expected proportion of immigrants in each patch at each generation, (*M*_m_ + *M*_f_)/(*n*_m_ + *n*_f_) (on a natural log scale, see Supplementary Figure 1 for linear scale;other parameters: *h*_f_ = *h*_m_ = 0.2, *M*_m_ = *M*_f_, *n*_m_ = *n*_f_ = 5, see Supplementary Figure 1 for effect of *F*-statistic in large groups). Bottom: selection according to ratio of homozygotic effects in females and males (*c*_m_/*c*_f_) and the expected proportion of males among immigrants in each patch at each generation (*M*_m_/(*M*_m_ + *M*_f_);other parameters: *n*_m_ = *n*_f_ = 5, *h*_f_ = *h*_m_ = 0.2, *M*_m_ + *M*_f_ = 0.1).

We also derived the equilibrium allele frequency when polymorphism is favoured under limited dispersal (see eq. A-47 in Appendix A.2.8). In line with eq. (5), we find limited dispersal increases the equilibrium frequency of the allele with the weaker detrimental effect in homozygotes, irrespective of dominance (e.g. if *c*_m_ < *c*_f_, the equilibrium frequency of the male-detrimental allele *A* increases with limited dispersal). In other words, where there is polymorphism, inbreeding leads selection to disproportionately direct equilibrium allele frequencies based on selective effects in homozygotes rather than heterozygotes.

#### 3.1.4 Non-additive effects: sex-biased dispersal changes the nature of sexually antagonistic polymorphism

Analysing eq. (5) further reveals that the nature of sexually antagonistic alleles maintained by balancing selection depends on immigrant sex ratio (Figure 3A bottom). Specifically, as dispersal becomes biased towards one sex, polymorphism tends to be favoured when the cost of the detrimental allele to the philopatric sex is greater than the cost of the detrimental allele to the dispersive sex (e.g. where dispersal is male-biased, *M*_m_/(*M*_m_ + *M*_f_) > 0.5, polymorphism is more likely when *c*_f_ > *c*_m_, see grey region Figure 3A bottom). Accordingly, sex-biased dispersal also relaxes the condition for fixation of the allele beneficial to the dispersing sex, and constrains the condition favouring fixation of the allele beneficial to the philopatric one (e.g. male-biased dispersal increases the parameter space for fixation of the male-beneficial allele but decreases parameter space for fixation of the female-beneficial allele, see blue and red regions in Figure 3A bottom). In line with our analysis of the additive case (eq. 3), these effects occur because sex-specific dispersal leads the philopatric sex to experience stronger kin competition, weakening the selective advantage of the allele favouring competitiveness in that sex relative to the other. As a result, sex-differences in dispersal proclivity lead to different types (i.e. competitiveness effects) of alleles being maintained under balancing selection, thereby influencing the nature of sexually antagonistic polymorphism.

#### 3.1.5 Effects of limited dispersal are similar at X-linked and autosomal loci

We also derived the frequency change of allele *A* when linked to an X-chromosome (our results also apply to Z-linkage in ZW species;see Appendix A.3.1-A.3.2 for derivations). Our analysis of the resulting allelic dynamics, which can be found in Appendix A.3.3, shows that the effects of limited dispersal on X-linked sexual antagonism are broadly the same as for autosomes. Specifically, inbreeding also leads to more constrained polymorphism conditions on the X-chromosome (Figure 3B top;see also Hitchcock and Gardner, 2020 for a recent discussion of inbreeding effects on X-linked sexual antagonism), and sex-specific dispersal also favours adaptation in the more dispersive sex at the expense of the other (so that polymorphism tends to be favoured when the cost of the detrimental allele is greater in the philopatric sex, Figure 3B bottom).

There are nevertheless some discrepancies between autosomal and X-linked positions due to the fact males only carry one copy of the X-chromosome. Firstly, the difference in copy number between the sexes means that male reproductive value for X-linked genes is half that of females (because on average, males transmit half as many X copies to their offspring as females). As a result, selection via female competition is twice as important as via male competition on the X-chromosome. The consequences of this can be most clearly seen by considering an X-linked additive locus (*h*_f_ = 1/2) with equal allelic effects in males and females (*c*_f_ = *c*_m_ = *c*).

At such a locus, we show in Appendix A.3.3 that the change in *p* per generation is given by

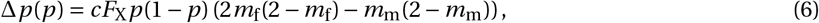

where,

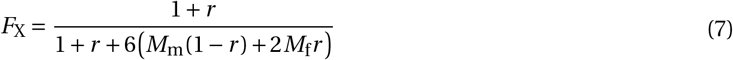

is the two-way coalescence probability for the X-chromosome (i.e. the probability two neutral genes sampled from the same sex in the same group before dispersal are IBD, in the limit of low dispersal and large group size, see Appendix A.3.2 for derivation of *F*_X_; note that eq. 7 is equivalent to eq. 9 in Ramachandran et al., 2008). Eq. (6) shows that under weak additive effects, selection favours the fixation of one allele or the other depending on sex-specific dispersal, like on autosomes. But unlike on autosomes, for the male beneficial allele to be favoured, males must be twice as likely to disperse than females (i.e. Δ*p* (*p*) < 0 for all 0 < *p* < 1 when *m*_m_ > 2 *m*_f_, by contrast this occurs when *m*_m_ > *m*_f_ on autosomes see eq. 3).

Comparing the probabilities that autosomal and X-linked genes are IBD (eq. 4 vs. eq. 7) reveals a second effect of male hemizygosity: that X-linked genes in individuals from the same group tend to coalesce faster than autosomal genes, with the exception of where sex and immigrant ratios are both strongly female biased (specifically, *F* > *F*_X_ only when *r* < 0.5 and *M*_m_/(*M*_m_ + *M*_f_) < 2*r* (2*r* − 1)/(8*r*^2^ − 5*r*-1), bottom left corner of Figure 2A). This is because there are always fewer X copies than autosomes in a group. Furthermore, like for autosomes, X-linked coalescence probability increases when the adult and immigrant sex ratios are biased in a similar manner, however this occurs to a greater extent on the X when both ratios are biased towards males (Figure 2B-C for exact probabilities). As a consequence, polymorphism conditions tend to be more restrictive on the X-chromosome than on autosomes with dispersal limitation (Figure 3A-B top), especially when immigrants are more likely to be males (Figure 3B bottom).

### 3.2 The interplay between sexually antagonistic selection and genetic drift under limited dispersal

By assuming an infinite number of groups, our analyses so far have ignored the influence of genetic drift, which is a pervasive evolutionary force in finite populations (Charlesworth and Charlesworth, 2010) and thought to be especially relevant for intralocus sexual conflict (Connallon and Clark, 2012; Mullon et al., 2012). Predicting how sexually antagonistic variation is maintained under the joint effects of selection and genetic drift in subdivided populations, however, is not straightforward. This is because dispersal limitation influences both balancing selection and genetic drift with conflicting implications for polymorphism. On one hand, because limited dispersal leads to kin competition and inbreeding, it weakens the strength of balancing selection relative to genetic drift. On the other, genetic isolation between patches increases the effective size (*N*_e_) of a population and therefore reduces the strength of genetic drift relative to selection (in the absence of patch extinction, Caballero, 1994; Wang and Caballero, 1999).

#### 3.2.1 A simulation approach

To investigate the conflicting effects of limited dispersal on the maintenance of polymorphism at sexually antagonistic loci through selection and genetic drift, we used individual-based simulations to track the dynamics of sexually antagonistic alleles at a di-allelic autosomal locus in a finite subdivided population (see Appendix B for full simulation procedure with SLiM 3.3, Haller and Messer, 2019). Our simulations follow a population of 50 groups each containing equal numbers of male and female adults (*n*_m_ = *n*_f_ = 5 for computational tractability, see also Supplementary Figure 2 for additional simulations where *n*_m_ = *n*_f_ = 10 that produced qualitatively equivalent results to those in this section).

At the beginning of a generation each female in a group produces a Poisson distributed number of eggs (mean *k* = 20). Males are then sampled at random with replacement within groups to fertilise each egg and produce a zygote. Zygotes are assigned male or female identity with equal probability (i.e. sex ratio is unbiased at birth). Male and female zygotes disperse to another randomly-selected group with sex-specific probability *m*_m_ and *m*_f_, or stay in their natal group otherwise. Finally, *n*_m_ and *n*_f_ males and female zygotes are sampled without replacement in each patch to reach adulthood, with the probability of a zygote being sampled weighted by the competitiveness of its genotype (according to Figure 1A).

We ran simulations for a number of dispersal regimes (*m*_m_,*m*_f_) and allelic effects (*c*_m_, *c*_f_). We also considered additive (*h*_m_ = *h*_f_ = 0.5) and dominance reversal (*h*_m_ = *h*_f_ = 0.2) scenarios. Simulations were started with equal frequencies of male- and female-beneficial alleles in the population (*p* = 0.5 at Hardy-Weinberg equilibrium), and then left to run until one allele is fixed and the other is lost (see green trajectories in Figure 4A for examples of runs). We ran 10,000 replicates for each parameter combination. For each simulation replicate, we tracked segregation time (number of generations until loss of an allele), the final frequency of the female-beneficial allele *A* (0 or 1), as well as the *F*_IT_ and *F*_ST_ statistics in adults (which were time-averaged across generations in a replicate, see purple and orange dots and lines in Figure 4A for example, and Appendix B for details).

**Figure 4:**
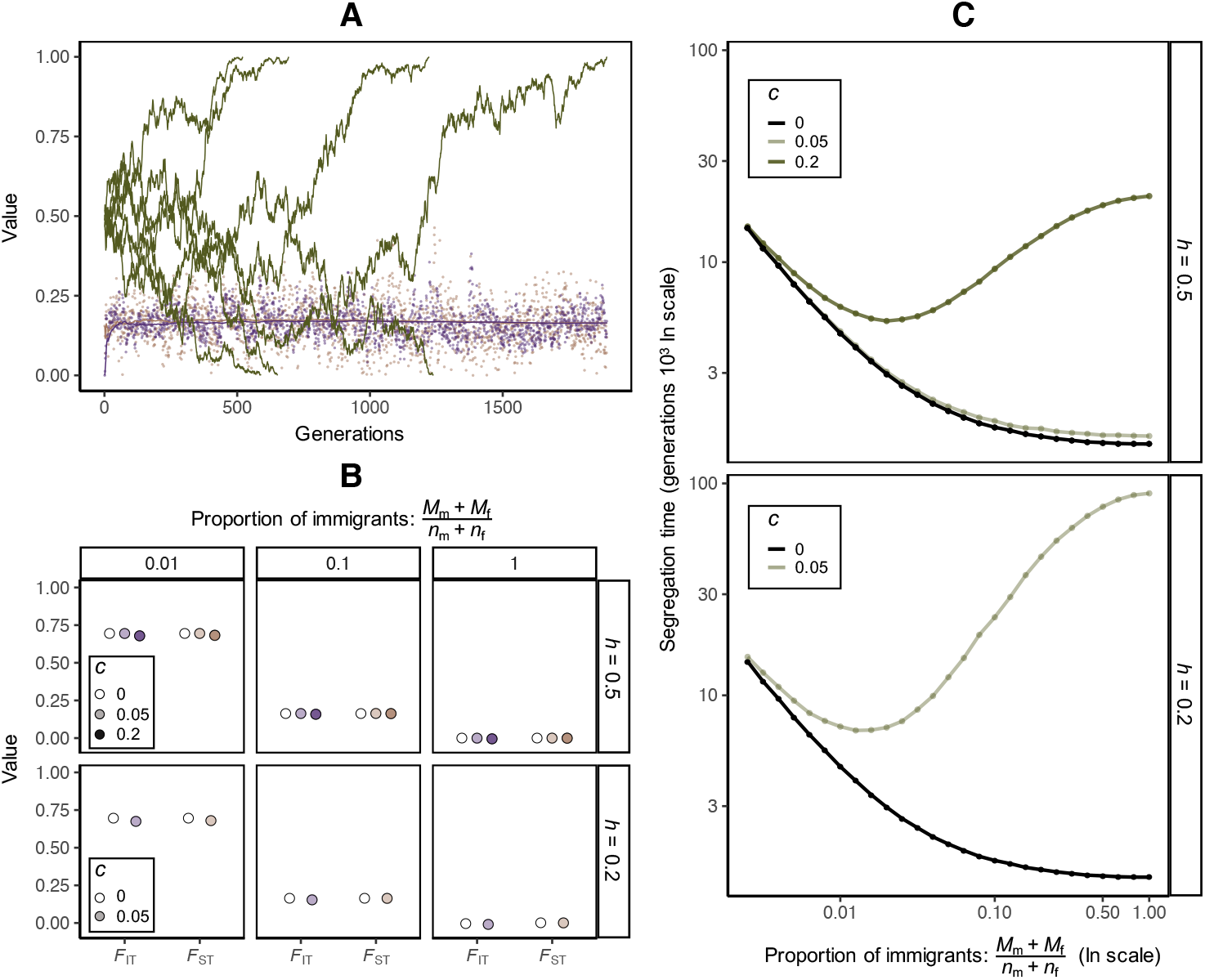
Allele segregation time and genetic variation in individual-based simulations. **A.** shows ex-ample simulation runs under neutrality (*c* = *c*_m_ = *c*_f_ = 0) and limited dispersal (when expected proportion of immigrants per group per generation, (*M*_m_ + *M*_f_)/(*n*_m_ + *n*_f_) = 0.1). Green trajectories show allele (*A*) frequency for different replicate runs, where the time taken for *A* to fix or be lost represents “segregation time” for that simulation. Purple and orange dots show *F*_IT_ and *F*_ST_ respectively at each generation during a simulation, while purple and orange lines represent time-averaged values for these quantities (see eqs. B-1 and B-2 in Appendix B for details on how *F*_IT_ and *F*_ST_ are calculated). **B.** shows time-averaged *F*_IT_ and *F*_ST_ averaged across all (10,000) replicate runs under different levels of limited dispersal (number at top of each column panel gives the expected proportion of immigrants in each group), dominance effects (row panels indicate *h* = *h*_f_ = *h*_m_), and strengths of sexually antagonistic selection (white circles represent neutrality *c* = 0, light and dark coloured represent *c* = 0.05 and *c* = 0.2 respectively). Note that because the *F*_ST_ and *F*_IT_ estimates for each replicate are already averages across generations in a simulation, these statistics are highly consistent between simulation runs (standard deviation of order 10^-2^ or lower). **C.** shows segregation time averaged across replicate runs as a function of total fraction of immigrants (which is on a natural log scale, see Supplementary Figure 2 for linear scale) and selection strength (black line for neutrality, solid light and dark green lines for *c* = 0.05 and *c* = 0.2 respectively, see Supplementary Figure 2 for *c* = 0.01), row panels again show different dominance effects. Dots refer to parameter (proportion of immigrants and strength of selection) combinations used in simulations. Other parameter values for simulations: *n*_m_ = *n*_f_ = 5, see Appendix B for more details on simulation procedure).

#### 3.2.2 Strong sexual antagonism, dominance reversal and frequent dispersal prolong polymorphism in finite populations

We first consider the case where allelic effects are equal in both sexes (*c* = *c*_m_ = *c*_f_) and where dispersal is unbiased (*m*_m_ = *m*_f_). In line with eq. (4), we observe that as dispersal probability decreases, inbreeding, *F*_IT_, and genetic differentiation among groups, *F*_ST_, both increase (irrespective of the strength of selection and dominance effects, Figure 4B). In other words, as dispersal becomes limited we see a reduction in genetic variation within individuals and within groups, as well as an increase in genetic variation between groups. Such conditions are known to increase effective population size (Wang and Caballero, 1999). Accordingly, limited dispersal prolongs the maintenance of neutral genetic variation in our simulations (*c* = 0, black line in Figure 4C).

For sexually antagonistic loci, the effect of dispersal on segregation time depends on whether sexual antagonism leads to balancing selection or not. Where alleles have weak additive (*h*_m_ = *h*_f_ = 0.5) effects that are symmetric across the sexes (*c*_m_ = *c*_f_ = *c* ≪ 1), selection is effectively neutral in the absence of sex-specific dispersal (eq. 3). In this case, the impact of limited dispersal on segregation time is largely similar to its effect on a neutral locus, with variation maintained for longer when dispersal is limited (compare black and pale green in Figure 4C top panel). By contrast, when selection is balancing, either due to dominance reversal (*h*_m_ < 0.5, *h*_f_ < 0.5, eq. 5) or strong selection (i.e., large *c*_m_ = *c*_f_ = *c*; see e.g., Fry, 2010, for a well-mixed population), we observe a quadratic relationship between segregation time and dispersal, with polymorphism maintained for longer when dispersal is either very strong or very weak (dark green curve in Figure 4C top panel and light green curve in Figure 4C bottom panel). Thus the balance between the opposing effects of population genetic structure on polymorphism, increasing *N*_e_ on one hand and decreasing the strength of selection on the other, varies with the degree to which total dispersal is limited.

#### 3.2.3 Sex-specific costs and dispersal interact to modulate segregation time

Finally, we consider the segregation of sexually antagonistic alleles where the allelic cost (*c*_f_ / *c*_m_) and immigrant (*M*_m_/(*M*_m_ + *M*_f_)) ratios can be biased, representing sex differences in selection strength and dispersal proclivity respectively. We assume dominance reversal (*h*_m_ = *h*_f_ = 0.2), weak selection (*c*_m_ + *c*_f_ = 0.1) and a fixed total proportion of immigrants (either (*M*_m_ + *M*_f_)/(*n*_m_ + *n*_f_) = 0.1 for moderate or (*M*_m_ + *M*_f_)/(*n*_m_ + *n*_f_) = 0.01 for strong dispersal limitation).

To understand the results of these simulations, it is useful to recall that sex-specific allele costs and dispersal together determine whether sexual antagonism generates balancing selection (Fig 3A bottom), as well as the equilibrium allele frequency, *p*\*, favoured by such selection (Figure 5A, see also Appendix A.2.8). When the cost and immigrant ratios are such that selection is balancing (so 0 < *p*\* < 1, Figure 5A), our simulations show that polymorphism is long-lived relative to neutral variation (grey, light blue and light red lines in Figure 5B). In particular, segregation time is greatest when *p*\* is close to 0.5 (i.e. furthest from the fixation boundaries, Figure 5A-B), in agreement with analyses of finite panmictic populations (Connallon and Clark, 2012; Mullon et al., 2012). However, the importance of these selective effects diminishes under strong dispersal limitation. Specifically, segregation time decreases relative to neutral variation, and is less sensitive to *p* *, when dispersal is strongly rather than moderately limited (compare left and right panels in Figure 5B). As before, this is because the strength of sexually antagonistic selection decreases relative to drift as dispersal become severely limited and kin competition increases.

**Figure 5:**
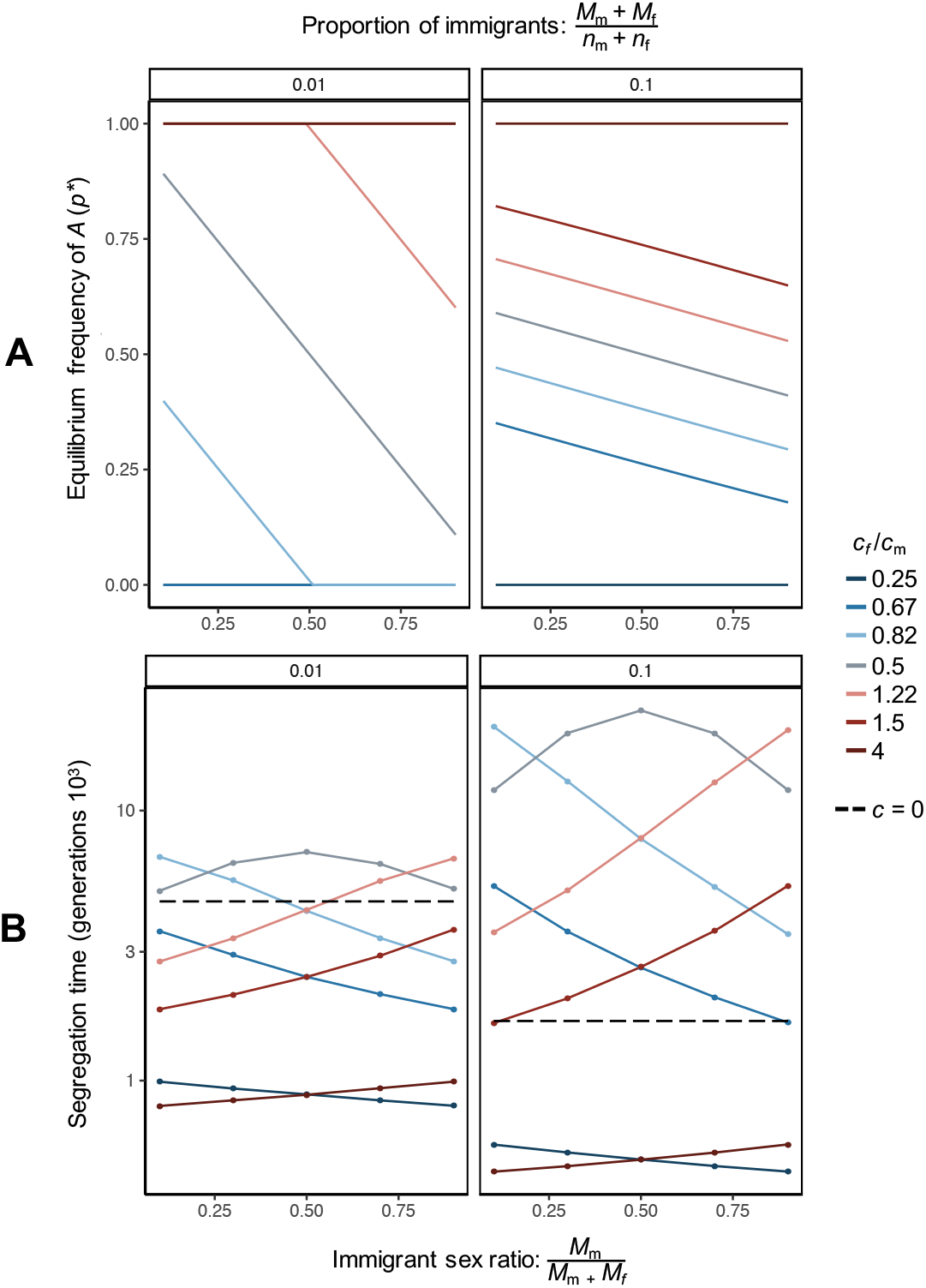
Fate of sexually antagonistic alleles according to sex-specific competitiveness and dispersal effects. Effects of selection and drift on sexually antagonistic alleles when allele cost (*c*_f_ / *c*_m_) and immigrant (*M*_m_/(*M*_m_ + *M*_f_)) sex ratios are varied. Blue and red lines represent male (*c*_f_ < *c*_m_) and female (*c*_f_ > *c*_m_)-biased competitiveness effects respectively, while grey represents equal costs in both sexes (*c*_f_ = *c*_m_). Left and right panels indicate different levels of dispersal limitation (Left: strong limitation with average proportion of immigrants per generation per group, (*M*_m_ + *M*_f_)/(*n*_m_ + *n*_f_) = 0.01;Right: weak limitation with (*M*_m_ + *M*_f_)/(*n*_m_ + *n*_f_) = 0.1). **A.** Equilibrium frequency of *A* under sexually antagonistic selection alone (i.e. assuming an infinite number of patches), calculated numerically by plugging exact coalescence probabilities (see Appendix A.2.7) into eqs. (1)-(2) and solving Δ*p*(*p*\*) = 0 for *p*\* (where selection is directional, we set *p*\* = 0or *p*\* = 1 depending on whether selection favours loss or fixation of *A*, respectively). Other parameters: *h*_f_ = *h*_m_ = 0.2, *n*_m_ = *n*_f_ = 5. **B.** Segregation time in individual-based simulations averaged across all replicate runs. Dots refer to parameter (cost and immigrant ratio) values used in simulations (see Appendix B for details). Other parameters: *h*_f_ = *h*_m_ = 0.2, *n*_m_ = *n*_f_ = 5, *c*_f_ + *c*_m_ = 0.1, dashed black line shows the neutral case (*c* = *c*_m_ = *c*_f_ = 0).

Where the cost and immigrant ratios produce directional selection (i.e. selection favours *p* = 0 or *p* = 1, Figure 5A), polymorphism is lost more rapidly at a sexually antagonistic than neutral locus (dark blue and light red lines in Figure 5B), with segregation time lowest when selection strength is most asymmetrical between the sexes, and when the dispersing sex is under the more intense selection. This is because these conditions provide the strongest directional selection and thus most rapid fixation of the favoured allele. Finally, contrary to where selection is balancing, strongly limited dispersal leads to an increase in segregation time for alleles under directional selection (left versus right panels in Figure 5B), as in this case heightened kin competition reduces the efficacy with which such selection purges variation.

## 4 Discussion

Our analyses indicate that limited dispersal can significantly impact evolution at loci under sexual conflict, altering both the nature and scope of sexually antagonistic variation through three main pathways. Firstly, limited dispersal leads to kin competition as individuals of the same group are more likely to carry identical alleles than random individuals. Such competition between relatives weakens the strength of selection on competitive traits and hence of sexually antagonistic balancing selection. In addition, where dispersal is sex-specific, the strength of kin competition is asymmetric between the sexes and consequently biases the intersexual tug-of-war over allele frequencies in favour of the more dispersive sex (Figure 3 Bottom). Secondly, limited dispersal causes inbreeding and thus reduced heterozygosity. This in turn limits the capacity of dominance reversal to promote polymorphism at sexually antagonistic loci (Figure 3 Top). Thirdly, in finite populations, limited dispersal simultaneously reduces both the strength of genetic drift and the strength of sexually antagonistic selection. The implications of this for the maintenance of polymorphism in smaller populations depend on the strength of sexual antagonism, with limited dispersal promoting long-term polymorphism when balancing selection is weak and impeding it when balancing selection is strong (Figure 4C). Our analyses indicate that these various effects of limited dispersal on the segregation of sexually antagonistic alleles are most relevant in subdivided populations characterized by *F*_ST_ values of order 10^-1^ or higher. Such values are consistent with many *F*_ST_ estimates for natural populations (e.g. mammals, Cegelski et al., 2003; Hammond et al., 2006; birds, Woxvold et al., 2006; Harrison et al., 2014; reptiles, Berry et al., 2005; Böhme et al., 2007; and amphibians Wang et al., 2020; Canestrelli et al., 2007 - see also p. 334 in Barton, 2001, p. 302 in Hartl and Clark, 2007 and p. 310 in Charlesworth and Charlesworth, 2010 for more general comments). These observations suggest that our model may be relevant for a wide range of taxa.

Our results lead to several predictions for interspecific variation in the distribution of sexually antagonistic alleles. Firstly, in populations with sex-specific dispersal, we expect that sexually antagonistic polymorphism is characterised by greater antagonistic costs, and thus greater fitness variation attributable to sexual antagonism, in the non-dispersing sex compared to the dispersing one (e.g. when males disperse more, polymorphism is expected at loci with *c*_f_ > *c*_m_). Although we are not aware of any empirical work explicitly investigating this, there is potentially relevant data from pedigree studies. In the yellow-pine chipmunk (*Tamias amoenus*), a species which shows stronger kinship between neighbouring females than males (implying male-biased dispersal, Dobson, 1982; Schulte-Hostedde et al., 2001), females are under directional selection for increased body size, while males are not (Schulte-Hostedde et al., 2002). This suggests that in this species, males are closer to their fitness optimum (so that alleles deviating from this optimum incur a small *c*_m_ in males) than females (so that genetic variation for body size in females cause a large *c*_f_). In the great reed warbler (*Acrocephalus arundinaceus*), which shows female-biased dispersal (Hansson et al., 2003), sexual antagonism has been described over wing length (Tarka et al., 2014). In line with our prediction, males are under stronger selection for wing length than females, and show larger additive genetic variance for fitness (Tables 1 and 2 in Tarka et al., 2014). However, the authors of this study also report a negative correlation between a male’s wing length and the fitness of his female relatives (Table 3 in Tarka et al., 2014) but not the reverse (a correlation between female wing length and the fitness of male relatives), leading them to suggest that alleles beneficial for males are in fact costlier to females. These conflicting results highlight the difficulty in interpreting the strength of sexually antagonistic allele effects from quantitative genetics studies. More generally, further work is needed to discern the extent to which these observations reflect wider trends, and we suggest future studies consider the effect of dispersal regimes and mating systems on relatedness when formulating hypotheses on sexually antagonistic variation.

Secondly, we expect long-term sexually antagonistic polymorphism to accrue less, relative to neutral variation, in dispersal limited populations than in well-mixed populations. Furthermore, in dispersal limited species with small total population size (e.g. small fragmented populations), we expect more sexually antagonistic variation than neutral variation when selection and dispersal are both strong. Broadly speaking, we therefore expect different levels of sexually antagonistic variation between species according to population genetic structure. However, because of the close link between dispersal, mating system and the strength of sex-specific selection (Hamilton, 1967; Greenwood, 1980; Shields, 1987; Perrin and Mazalov, 2000; Li and Holman, 2018), taxa with contrasting spatial structures may also show differences in intensity of intralocus conflict, confounding interspecific patterns in sexually antagonistic variation. Instead, a more fine-scale test would be provided by a species with multiple populations of differing spatial structure but similar levels of sexual antagonism. One such potential study system is the Atlantic salmon (*Salmo salar*), a species lacking sex chromosomes in which QTL analyses have identified a polymorphic sexually antagonistic locus that is maintained through sex-specific dominance (Figure 1D, Barson et al., 2015). Different salmon populations exhibit different levels of structure, with Scandinavian populations showing strong subdivision (Vähä et al., 2007, 2008) but North American populations appearing relatively panmictic (Wellband et al., 2019). Under the assumption that both salmon populations experience similar patterns of sexually antagonistic selection, this species would be well suited to test the effects of spatial subdivision on the amount of sexually antagonistic variation maintained in a population.

Another, more broad-brush, prediction from our results arises from the observation that sex-specific selection over reproductive success tends to be stronger in males (Janicke et al., 2016; Singh and Punzalan, 2018). In the context of our study, stronger male selection should cause sexually antagonistic alleles to have greater costs in males than females (i.e. *c*_m_ > *c*_f_ in our notation). Where this is true, our results suggest that sexually antagonistic polymorphism is more likely in taxa showing male philopatry (i.e. female-biased dispersal), as heightened male kin competition cancels out the effects of stronger selection for male reproductive success potentially leading to balancing selection (Fig 3 bottom). Conversely, in taxa where dispersal is predominantly male-biased, the combined effects of sexual antagonism and kin competition in females should favour the fixation of male-beneficial alleles, leading to less sexually antagonistic variation and male adaptation at the expense of females.

In addition to being useful to understand between-species and between-population patterns of sexually antagonistic variation, our results also have implications for variation within genomes. Much debate in studies of sexual antagonism has centred around the likelihood of different genomic regions accumulating sexually antagonistic polymorphisms, in particular whether such variation should be more common on the X-chromosome or autosomes (Rice, 1984; Gibson et al., 2002; Gavrilets and Rice, 2006; Fry, 2010; Otto et al., 2011; Mullon et al., 2012; Jordan and Charlesworth, 2012; Ruzicka and Connallon, 2020). Previous results have pointed out potential disparities between the X-chromosome and autosomes in their proclivity to harbour sexually antagonistic variation due to male hemizygosity for the X, which leads to differences in selection (i.e. dominance effects act only through females for X-linked genes, Rice, 1984; Gavrilets and Rice, 2006; Fry, 2010; Jordan and Charlesworth, 2012), and differences in the strength of genetic drift (i.e. stronger genetic drift on the X owing to fewer copies of X-chromosomes than autosomes, Charlesworth et al., 1987; Vicoso and Charlesworth, 2009; Mullon et al., 2012). Our findings indicate that population subdivision influences both of these differences. First, limited dispersal diminishes the importance of selection on heterozygotes and thus dominance effects in driving differences in variation between X-linked and autosomal locations (Figure 3). Second, sex-specific dispersal and local demography alter *F*_ST_ for X- and autosomally-linked genes in different ways (Figure 2) and hence influence the effective population size of the X-chromosome relative to an autosome. In particular, the X-linked and autosomal effective population sizes converge if the adult sex ratio within patches is sufficiently female-biased under limited dispersal (when *r* ≈ 0.15, see p. 296 in Ramachandran et al., 2008). Taken together, these results suggest that conditions under which the X-chromosomes and autosomes differ in their propensity to retain sexually antagonistic alleles are less straightforward than predicted by panmictic models. Thus, limited dispersal may help explain the current patchy empirical support for either chromosome region harbouring disproportionate levels of sexually antagonistic variation (e.g. Gibson et al., 2002; Lucotte et al., 2016; Grieshop and Arnqvist, 2018; Ruzicka et al., 2019).

Our analysis highlights how limited dispersal impacts sexual antagonism by introducing genetic correlations within individuals and groups (via *F*_IT_ and *F*_ST_), skewing genotype frequencies away from Hardy-Weinberg proportions. Alongside limited dispersal, other factors can distort allele distributions away from Hardy Weinberg equilibrium and thus influence selection on sexually antagonistic variation. In fact, previous theoretical studies have shown that by increasing and decreasing the level of inbreeding respectively, self-fertilisation (Jordan and Connallon, 2014; Tazzyman and Abbott, 2015) and assortative mating for fitness (Arnqvist, 2011; Kasimatis et al., 2019; Hitchcock and Gardner, 2020) affect the potential for sexually antagonistic polymorphism (see also Kasimatis et al., 2019 for assortative mating for genotype). Meanwhile, *F*_ST_ can be influenced by life-history traits other than dispersal, such as iteroparity (which increases genetic relatedness within groups, Taylor and Irwin, 2000). Interestingly, sex-asymmetries in relatedness also arise in systems where the sexes differ in the extent to which they are itero- or semelparous (e.g. one sex is long-lived while the other is short-lived, Van Cleve et al., 2010), suggesting that sex-differences in life-history strategies may have similar consequences for sexual antagonism as sex-specific dispersal.

Although our study was framed in the context of sexual antagonism, we also expect population subdivision to have implications for other forms of antagonistic evolution, such as antagonistic pleiotropy (where alleles have antagonistic effects on different fitness components, e.g. Williams, 1957; Rose, 1982; Kirkwood and Rose, 1991; Curtsinger et al., 1994; Williams and Day, 2003; Connallon and Clark, 2012; see also Zajitschek and Connallon, 2017, 2018, for antagonistic pleiotropy in the context of sexual antagonism under panmixia). Here, models have also highlighted the importance of component-specific dominance effects, such that antagonistic effects are recessive for each fitness component, in facilitating balancing selection (Rose, 1982, 1985; Curtsinger et al., 1994; Connallon and Chenoweth, 2019). But these models assume populations are well-mixed and that genotype frequencies prior to antagonistic selection are at Hardy-Weinberg equilibrium (Rose, 1982, 1985; Curtsinger et al., 1994). From our results, we expect that limited dispersal will also reduce the scope for polymorphism from antagonistic pleiotropy and may therefore influence the evolution of a wide range of traits involved in fitness trade-offs (Roff, 2002), such as viability and expression of secondary sexual characters (Zahavi, 1975; Lande, 1981; Kirkpatrick, 1982; Grafen, 1990; Iwasa et al., 1991; Iwasa and Pomiankowski, 1994), resource acquisition (Van Noordwijk and de Jong, 1986), and reproductive investment and lifespan (Williams, 1957; Williams and Day, 2003; Nussey et al., 2013).

Our model, which has allowed us to produce tractable and intuitive results about intralocus sexual conflict under limited dispersal, relies on several assumptions that are of course unrealistic for many natural populations. In particular, while our model allows us to vary demographic parameters such as adult sex ratio and dispersal regime (immigrant sex ratio and total dispersal rate), these quantities are not free to vary among groups or evolve. In natural populations, however, parameters such as sex ratio and sex-specific dispersal are correlated by ecological dynamics, and their interdependence will lead them to co-evolve. Investigating how such “eco-evo feedbacks” between local demography and sexual antagonism shape mating system evolution would be an interesting avenue for future work (Svensson, 2019; Giery and Layman, 2019, for general considerations; see de Vries and Caswell, 2019 for such an “eco-evo” model of sexual antagonism under panmixia;and Rousset and Ronce, 2004 or Mullon and Lehmann, 2018 for analyses of selection on traits influencing demography in subdivided populations). In this context, it would also be relevant to relax our assumption that female fecundity is large in order to consider the effects of sex-specific reproductive variance (Mullon et al., 2014), which is common in nature (Clutton-Brock, 2007) and is especially important for selection in subdivided populations (Lehmann and Balloux, 2007). Another assumption of our model is environmental homogeneity between patches (that selection on males and females is consistent across space). In contrast to our results, a previous model has shown that in the absence of genetic structure (i.e. assuming infinite patch size), sex-biased dispersal in spatially heterogeneous environments can lead sexually antagonistic selection to favour adaptation in the philopatric sex over the dispersing one (Connallon et al., 2019). This is because under spatial heterogeneity, the dispersing sex experiences greater variance in selection pressure, weakening the overall strength of directional selection in this sex. To better understand the contrasting effects of spatial variation and genetic structure, a valuable extension of our model would therefore be to consider heterogeneous subdivided populations consisting of finite groups (although note this is significantly more complicated mathematically, e.g. Rousset and Ronce, 2004; Lehmann et al., 2016).

In conclusion, through its effects on inbreeding, kin competition and genetic drift, limited dispersal qualitatively alters the segregation of sexually antagonistic alleles, narrowing the scope for sexually antagonistic polymorphism and changing the nature of intralocus conflict. Dispersal patterns, and in particular their sex-specificity, are therefore a relevant consideration for the genetic and ecological conditions expected to lead to sexually antagonistic variation, as well as the nature of the alleles underlying such variation. More broadly, our results reinforce the general notion that spatial demography and population structure are important factors for population genetic dynamics and the maintenance of non-neutral variation (e.g. Wright, 1949; Hamilton, 1964; Hartl and Clark, 2007; Rousset, 2004; Charlesworth and Charlesworth, 2010).

## Acknowledgements

We thank Carl Mackintosh, Max Reuter, Stu Wigby, Henry Barton, Daniel Bolnick, Stephen Chenoweth and two anonymous reviewers for comments and discussion on a previous version of the manuscript, Ben Haller for advice on simulations, and the UK Natural Environment Research Council and the Swiss National Science Foundation (PCEFP3181243 to CM) for funding.

## Author contributions

EOF, VS and CM conceived the study. EOF and CM performed the analyses. EOF and CM wrote the manuscript with input from VS.

## Appendix

## A Allele frequency change in a subdivided dioecious population

In this appendix, we derive the change in frequency over one generation for an allele at an autosomal and sex-linked locus, assuming that selection is weak. Our derivation follows closely previous analyses of population genetics models in subidvided populations (Roze and Rousset, 2003; Rousset, 2004, in particular Van Cleve et al., 2010).

### A.1 Model

We first provide further details on our modelling assumptions, which are necessary for our derivations. Recall we consider a dioecious population that is divided among a large number *n*_g_ of groups, each composed of *n*_m_ and *n*_f_ adult males and females, respectively. The life-cycle of this population is as follows. 1) Adults mate randomly within their group, females then produce a large number *k* of male and female juveniles in equal proportions (i.e. unbiased sex ratio at birth). 2) Independently of one another, each juvenile either remains in its natal group, or disperses to another randomly chosen group. 3) Adults die, and male and female juveniles compete to fill the *n*_m_ and *n*_f_ open breeding spots in each group (so competition occurs within the sexes and within groups).

Each individual carries a quantitative phenotype that is genetically-encoded and that determines the competitiveness of a juvenile to settle in a group (step 3 of the life-cycle). We will separately consider evolution at two loci that influence this phenotype: at an autosomal and at a sex-linked locus (on the X-chromosome in an XY species or the Z chromosome in a ZW species). In either case, two alleles *A* and *a* with sexually antagonistic effects segregate in the population. We are interested in deriving the change in frequency in the whole population of the allele *A*. For mathematical convenience, the population is censused in juveniles before dispersal (between steps 1 and 2 of the life-cycle).

### A.2 Autosomal locus

#### A.2.1 Genotypes

To denote the genotype of males and females, we let 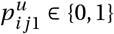 and 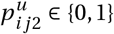 be the frequency of *A* at the maternally and paternally inherited locus, respectively, of juvenile *j* ∈ {1,…, *n*_f_*k*/2} of sex *u* ∈ {m, f} residing in group *i* ∈ {1,…, *n*_g_} (recall that *k* is the total fecundity of a female, so that within a group there are *n*_f_*k*/2 male and *n*_f_*k*/2 female juveniles after reproduction and before dispersal). The frequency of *A* in such an individual then is

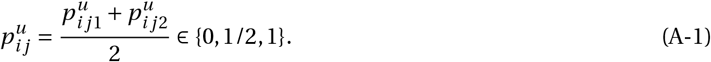

For example, if male 2 in group 1 has inherited *A* from its mother and *a* from its father, then 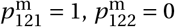, and 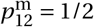. The average frequency of *A* in individuals of sex *u* can then be defined as

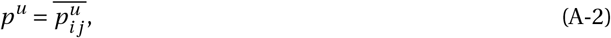

where the upper bar means the average over all juveniles *j* and over all groups *i* throughout.

#### A.2.2 Phenotypes

The evolving phenotype determines the competitiveness of male and female juveniles. We assume that baseline competitiveness is 1 and that allele *A* is detrimental to males (causing a decrease in competitiveness *δc*_m_ in homozygotes and *δh*_m_*c*_m_ in heterozygotes, Table A-1). Conversely, allele *a* is detrimental to females causing a decrease in competitiveness *δc*_f_ in homozygotes and *δh*_f_*c*_f_ in heterozygotes, Table A-1). Parameters *c*_m_ > 0 and *c*_f_ > 0 respectively capture the cost of the detrimental allele in male and female homozygotes (*δ* > 0 tunes the strength of selection), while parameters *h*_m_ and *h*_f_ determine the dominance or penetrance of the detrimental allele in males and females, respectively. We can then write the phenotype 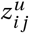 of juvenile *j* of sex *u* that resides in group *i* in terms of the frequency of allele *A* in this individual as

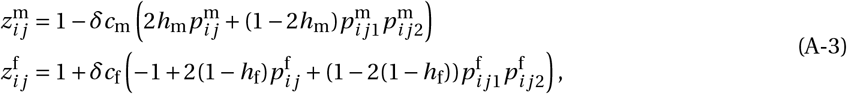

according to Table A-1.

**Table A-1.** Male and female phenotypes according to genotype.

|  | *aa* | *Aa* | *AA* |
| --- | --- | --- | --- |
| male phenotype | 1 | $1 - \delta h_m c_m$ | $1 - \delta c_m$ |
| female phenotype | $1 - \delta c_f$ | $1 - \delta h_f c_f$ | 1 |

#### A.2.3 Weighted allele frequency change

To track the dynamics of the frequencies of the *A* allele in the population, we consider the dynamics of a reproductive-value weighted average *p* of male and female frequencies,

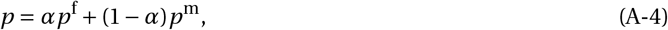

where *α* and 1 − *α* are the female and male reproductive values, respectively (e.g. Roze and Rousset, 2003). This weighting ensures that in the absence of selection (when *δ* = 0), the change in *p* is zero. For an autosomal locus, males and females have equal reproductive values: *α* = 1/2.

Following Roze and Rousset (2003) (adapting their eq. 35 for an autosomal locus) and Van Cleve et al. (2010) (their eq. 2), the change over one iteration of the life-cycle in weighted allele frequency change is

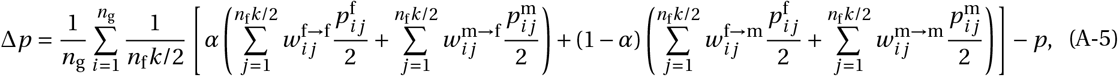

where 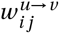 is the expected number of juveniles of sex *v* produced by juvenile *j* of sex *u* in group *i* (before dispersal) and the division of individual gene frequencies by 2 reflects that a gene has a probability of 1/2 to be passed to an offspring at an autosomal locus (with no segregation bias). There are thus four relevant fitness functions: 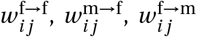, and 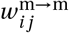. We consider these fitness functions in more detail later (section A.2.5). But first note that when sex ratio at birth is equal, a juvenile of either sex is expected to produce the same numbers of male and female juvenile so that 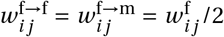 and 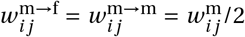, where 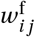 and 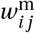 are the expected total numbers of juveniles produces by a female and male juveniles, respectively. Substituting for these total fitness expressions and for reproductive value *α* = 1/2 into eq. (A-5), we obtain

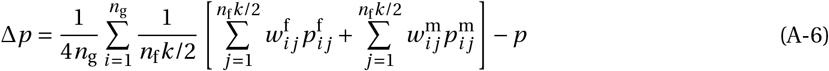

for the change in weighted allele frequency.

#### A.2.4 Selection gradient

Next, we connect individual fitness with individual gene frequencies. To do so, first note that since competition is within groups and between individuals of the same sex, male and female fitness functions can in general be written as functions of three relevant phenotypic values,

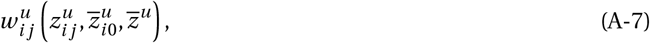

where 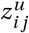 is the phenotype of the focal individual whose fitness is being considered (indexed *j* of sex *u* in group *i*); 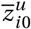 is the average phenotype among the other individuals of the same sex born in group *i* (i.e. excluding the focal individual); and 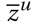 is the average phenotype of individuals of sex *u* in the population (see eq. A-30 for an example of such fitness function). Second, we use the chain rule on these fitness functions to obtain,

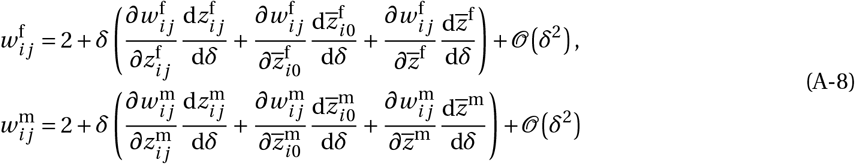

where 2 is the expected total number of juveniles produced by an individual under neutrality (i.e. one male and one female juvenile when *δ* = 0) and all derivatives are evaluated at *δ* = 0 (so at 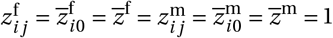, from eq. A-3). The above equation may be further simplified by using the following relationships

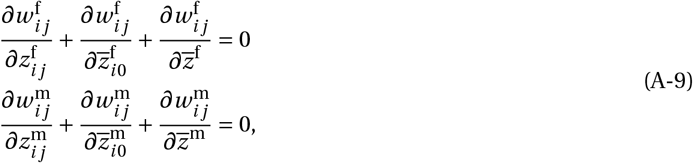

which hold because the population of each sex is constant, so that any fitness gain made by a focal individual due to a change of phenotype must be compensated by a decrease in fitness by the rest of the population (Rousset, 2004, p. 96).

Substituting eq. (A-9) into eq. (A-8), we obtain

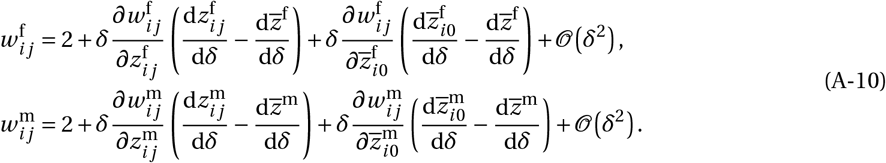

In turn, plugging eq. (A-10) into eq. (A-6), we obtain that the allele frequency change is

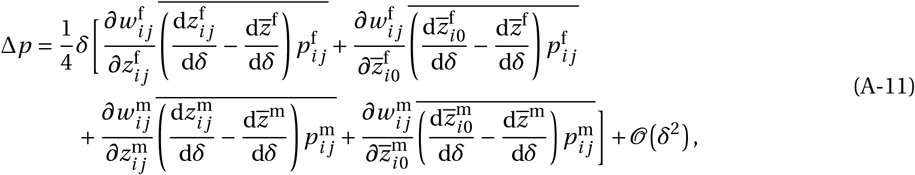

where the upper bar means the average over all juveniles and over all groups. From eq. (A-3), the averages in eq. (A-11) can be expressed in terms of products of allele frequencies within and between individuals:

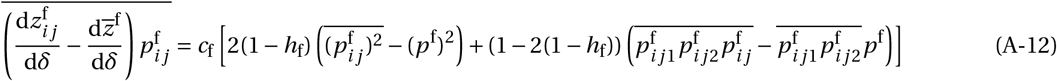

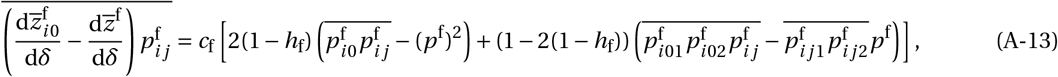

in females, where 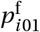 (and 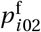) is the average frequency of the *A* allele at the maternal (and paternal) locus of the neighbouring females of a focal female indexed *j* in group *i*, and 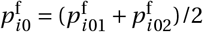. Similarly,

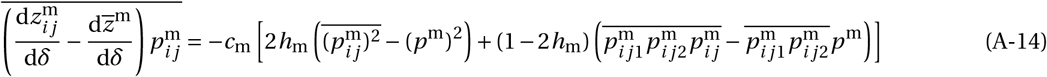

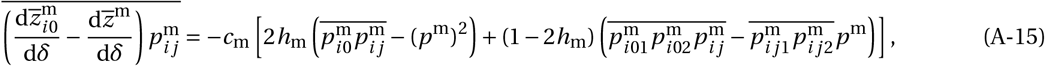

in males, where 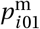 (and 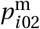) is the average frequency of the *A* allele at the maternal (and paternal) locus of the neighbouring males of a focal male indexed *j* in group *i*, and 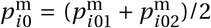. We first simplify eqs. (A-12) and (A-14). Using the properties of indicator variables (that 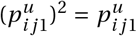 and 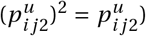, we have

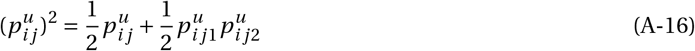

so that

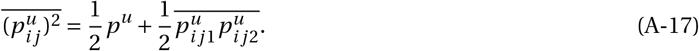

Similarly, we have that

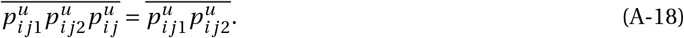

Substituting eqs. (A-17) and (A-18) into eqs. (A-12) and (A-14), these read as

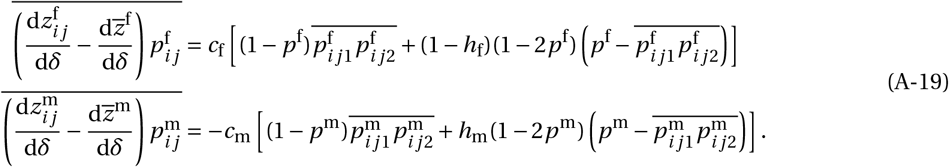

Next, we use the fact that the different averages in eqs. (A-13), (A-15), and (A-19) correspond to different probabilities of genetic identity (Roze and Rousset, 2003; Rousset, 2004; Van Cleve et al., 2010). For example, 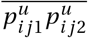 corresponds to the probability that two homologous gene copies randomly sampled in a juvenile of sex *u* before dispersal are both *A*. Further, these probabilities only need to be computed under neutrality to determine allele frequency change to the first order of *δ* (as they are already multiplied by *δ* in eqs. A-11). In such circumstances, the probabilities of identity can be connected with probabilities of identity-by-descent of neutral genes, which are independent of the frequency of *A* in the infinite island model (see Roze and Rousset, 2003 for further considerations). Take 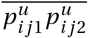 again for example. Consider the two lineages of these genes backward in time: either the two lineages have stayed in the same group and coalesced (with a probability that we denote *F*_IT_ and which we will show later is equal in males and females, see eq. A-41) or have dispersed to different groups patches and have not coalesced (with probability 1 − *F*_IT_). The probability that these two genes are *A* is then equal to *F*_IT_*p^u^* + (1 − *F*_IT_)(*p^u^*)^2^, where *p^u^* is the frequency of *A* in the population of individuals of sex *u*. However, this frequency is equal in males and females under neutrality (*p*^m^ = *p*^f^ = *p*) so that we have

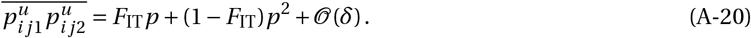

The probability *F*_IT_ is thus equivalent to Wright’s coefficient of inbreeding in the infinite island model. We will later compute this coefficient explicitly in terms of demographic parameters in section A.2.7. First, we express the other two averages featuring in eqs. (A-13) and (A-15) (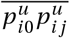 and 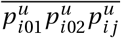) in terms of coalescence probabilities.

The probability, 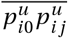, that two genes randomly sampled in different juveniles of the same sex *u* within the same group are both *A* can be expressed as

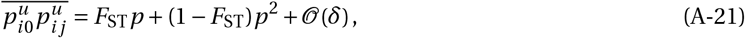

where *F*_ST_ is the probability that the two lineages of our two genes have stayed in the same group and coalesced (which is equal when those two genes are sampled in males and in females, see eq. A-36 for more details). This probability is equal to Wright’s *F*_ST_ within sexes in the infinite island model.

Finally, we need 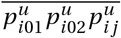, which corresponds to the probability that the two homologous genes of a juvenile of sex *u* and a third gene sampled in another juvenile of the same sex from the same group are all *A*. To compute this quantity, we let *K* be the probability that the three lineages of these genes have stayed in the same group and coalesced; and *L* be the probability that exactly two of those have coalesced (both probabilities are insensitive as to whether genes are sampled in males or females, see eq. A-43). With this notation, we have

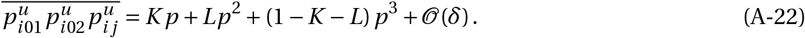

Further, from the relationships between the different coalescence events, we can write the probability *L* that exactly two of the genes of interest have coalesced as *L* = *F*_IT_ − *K* + 2(*F*_ST_ − *K*), where *F*_IT_ − *K* is the probability that the two homologous gene copies coalesce together but not with the gene sampled in another juvenile, and 2(*F*_ST_ − *K*) is the probability that the gene sampled in another juvenile and one gene from the homologous copies coalesce but not with the other copy. Substituting for *L* into eq. (A-22), we obtain

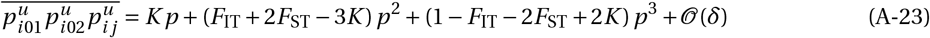

for the probability that the two homologous genes of a juvenile of sex *u* and a third gene sampled in another juvenile of the same sex from the same group are all *A*.

Plugging eqs. (A-20), (A-21) and (A-23) into eqs. (A-13), (A-15) and (A-19), which are in turn substituted into eq. (A-11), we get that the change in allele frequency is given by

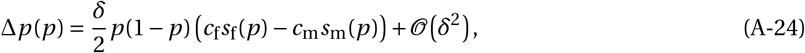

where

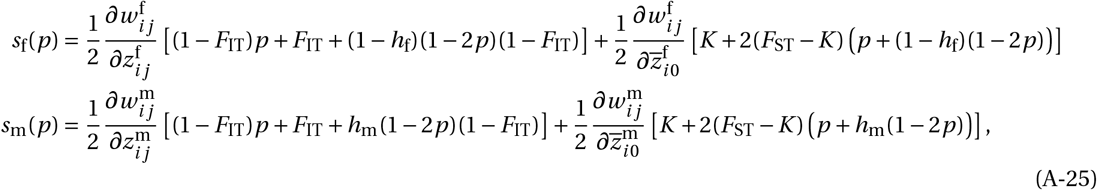

capture selection on the *A* allele due to its effect in females and males, respectively. We proceed to specify these selection effects in terms of demographic parameters and sex-specific dispersal, by specifying first the relevant fitness effects (i.e. fitness derivatives in section A.2.5), and second, relevant coalescence probabilities (in section A.2.7).

#### A.2.5 Fitness

According to the life-cycle described in section A.1, the expected number of female and male juveniles produced by a focal female juvenile can be decomposed as

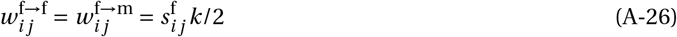

where 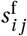. is the probability that this female juvenile survives competition to adulthood and *k*/2 is the number of male and female juveniles produced by an adult female. When the number of groups *n*_g_ is large and selection strength *δ* is small, the survival probability of a focal female is related to its phenotype 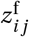. (i.e. its competitiveness) according to

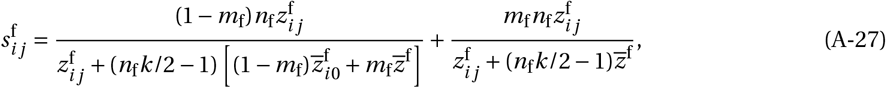

where *m*_f_ is the probability that a female juvenile disperses; 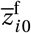 is the average phenotype among the other female juveniles born in the focal group (i.e. excluding the focal female); and 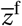 is the average female phenotype in the population. The first summand of eq. (A-27) is the probability that the focal female remains in its natal group (with probability 1 − *m*_f_) and survives, and the second, that she disperses (with probability *m*_f_) and survives. In both cases, her survival depends on the ratio of her competitiveness 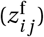 and the expected average competitiveness in the group she is in.

Similarly, the expected number of male and female juveniles produced by a focal male juvenile is

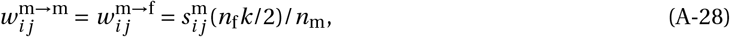

where 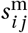 is the probability that this male juvenile survives to adulthood and (*n*_f_*k*/2)/*n*_m_ is the number of male and female juveniles produced by an adult male under random mating. Mirroring eq. (A-27), the survival probability of a focal male with phenotype 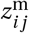 can be written as

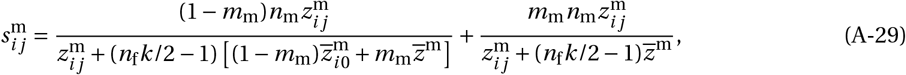

where *m*_m_ is the probability that a male juvenile disperses; 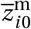 is the average phenotype among the other male juveniles born in the focal group; and 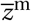 is the average male phenotype in the population.

Substituting eq. (A-27) into eq. (A-26) and eq. (A-29) into eq. (A-28), we obtain that total female, 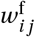., and male, 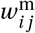, fitness (the expected total number of juveniles produced by a focal female and male juvenile) are given by

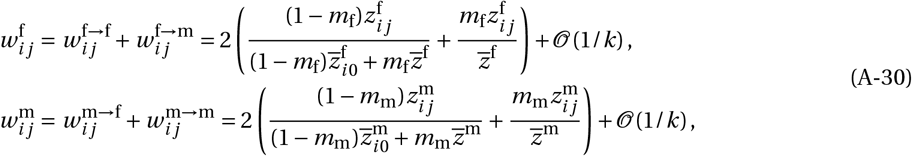

when fecundity *k* is large, which we will henceforth assume. From eq. (A-30), the direct and indirect fitness effects of female and male competitiveness are respectively given by

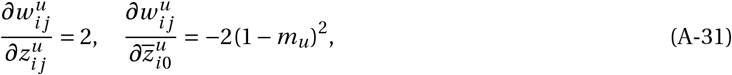

for *u* ∈ {m,f}. Plugging eq. (A-31) into eq. (A-25), we finally obtain main text equations (2).

#### A.2.6 Soft Selection

Eq. A-31 is based on the assumption that dispersal precedes selection, i.e. “hard selection”. Where dispersal occurs after selection (“soft selection”, Roze and Rousset, 2003; Débarre and Gandon, 2011), the survival of a focal juvenile depends only on its own phenotype and the phenotypes of individuals born in the same patch, specifically

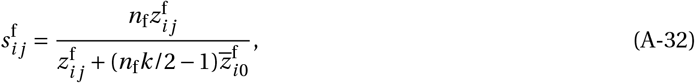

and

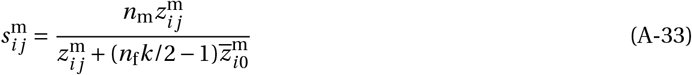

(compare with eqs. A-27 and A-29).

By plugging eq. (A-32) into eq. (A-26) and eq. (A-33) into eq. (A-28), and assuming large *k*, we find the direct and indirect fitness effects of competitiveness under soft selection are simply

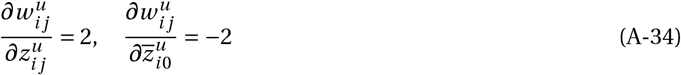

for *u* ∈ {*m, f*}. Substituting eq. (A-34) into eq. (A-25), we get an alternative version of main text equations (2),

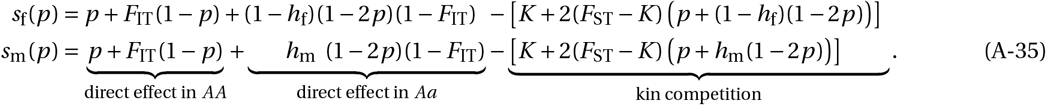

From eqs. (A-35) we can see that under soft selection, limited dispersal will narrow the scope for sexually antagonistic polymorphism by restricting the conditions where balancing selection is favoured (as under hard selection, main text eq. 5). However, unlike the hard selection case, the strength of kin competition is not sexspecific here (i.e. the kin competition term is not discounted by the probability of philopatry in each sex, as in main text eqs. 2). This is because selection occurs prior to dispersal so that all individuals experience kin competition in proportion to the average level of relatedness in their natal patch, which is equal between the sexes due to random mating within groups. More broadly, this means that sex-biased dispersal does not change the nature of polymorphism under soft selection (in contrast to hard selection where competitiveness is more strongly favoured in the dispersing sex, see Figure 3A bottom).

#### A.2.7 Coalescence probabilities at a neutral autosomal locus

In this section, we specify coalescence probabilities relevant for selection eq. (A-25) in terms of demographic parameters (numbers of males and females, *n*_m_, *n*_f_, and sex-specific dispersal, *m*_m_, *M*_f_). We do so using standard identity-by-descent arguments (e.g., Wang, 1997; Roze and Rousset, 2003; Rousset, 2004; Ramachandran et al., 2008; Van Cleve et al., 2010).

##### Pairwise coalescence probabilities

Consider first, *F*_ST_(*t*+1), the probability that two neutral genes randomly sampled at some arbitrary generation *t*+1 in different juveniles of the same sex *u* within the same group (before dispersal) coalesce. With probability 1/4 both sampled genes are maternally inherited; with probability 1/4, they are paternally inherited;and with probability 1/2, one is maternally inherited and the other paternally inherited. We can then write the probability that these two genes coalesce as

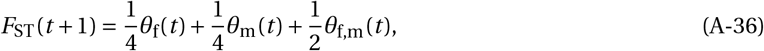

where *θ*_f_(*t*) and *θ*_m_(*t*) are respectively the probabilities that two maternally and paternally inherited genes coalesce; and *θ*_f,m_(*t*) the probability that one paternal gene and one maternal gene (in different individuals) coalesce. Consider the first of these, *θ*_f_(*t*), the probability that two maternally inherited genes coalesce. This probability will depend on whether or not the two juveniles that carry the sampled maternal genes have the same mother: 1) If the focal juveniles have the same mother (which occurs with probability 1/*n*_f_), then with probability 1/2 the two genes are the same copy and they coalesce with probability 1, or, with probability 1/2, the two genes are the homologous copies in the mother, in which case they coalesce with probability *F*_IT_(*t*) (the probability that two homologous gene copies in a juvenile sampled at generation *t* coalesce); 2) If the focal juveniles have different mothers (which occurs with probability 1 − 1/*n*_f_), then in order for the genes to coalesce both mothers must have remained philopatric (with probability (1 − *m*_f_)^2^) and their genes coalesce with probability *F*_ST_(*t*). From these considerations, we thus have

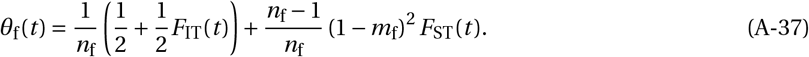

Similarly, the probability that two paternally inherited genes coalesce is

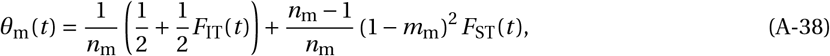

where 1/*n*_m_ is the probability that two juveniles have the same father. Finally, for one paternal gene and one maternal gene to coalesce, both the mother and father of the relevant juveniles must have remained philopatric, so that

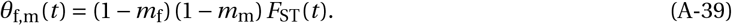

Substituting eqs. (A-37)-(A-39) into eq. (A-36), we obtain a recurrence for *F*_ST_(*t*),

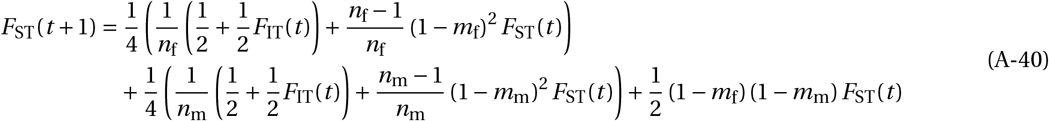

that also depends on *F*_IT_(*t*), the probability that two homologous gene copies in a juvenile sampled at generation *t* coalesce. This probability, meanwhile, satisfies the recurrence

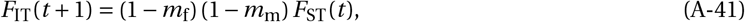

as for the two homologous gene copies of a juvenile to coalesce, both its mother and father must have remained philopatric, in which case the coalesce with probability *F*_ST_(*t*).

Solving for the equilibrium, *F*_ST_(*t* + 1) = *F*_ST_(*t*) = *F*_ST_ and *F*_IT_(*t* + 1) = *F*_ST_(*t* + 1) = *F*_IT_, using eqs. (A-40)-(A-41), we obtain

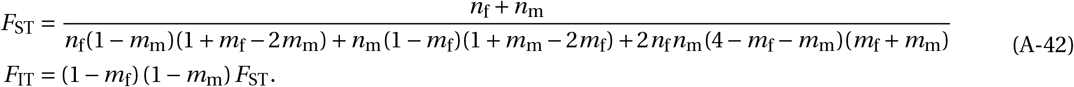

Substituting for *m*_f_ = *M*_f_/*n*_f_, *m*_m_ = *M*_m_/*n*_m_, *n*_f_ = (1 − *r*)*n* and *n*_m_ = *rn* in the above equation, we obtain *F* in eq. (4) of the main text in the limit *n* → ∞ (which is equivalent to e.g. eq. 8 of Ramachandran et al., 2008).

##### Threeway coalescence probabilities

To compute the probability, *K*, that the two homologous genes of a juvenile (which we label as juvenile “ 1” for the sake of argument) and a third gene sampled in another juvenile (labelled juvenile “2”) of the same sex sampled from the same group before dispersal all coalesce, consider that for this to happen, it is necessary that both parents of juvenile 1 are philopatric, which occurs with probability (1 − *m*_f_) (1 − *m*_m_). Then, with probability 1/2, the gene sampled in juvenile 2 is maternally inherited, in which case all three genes coalesce with a probability we call Θ_f_(*t*); with probability 1/2, the gene sampled in juvenile 2 is paternally inherited, in which case all three genes coalesce with a probability we call Θ_m_(*t*), so that

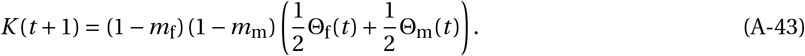

We proceed to specify Θ_f_(*t*) and Θ_m_(*t*).

Consider Θ_f_(*t*) first. If the gene sampled in juvenile 2 is maternally inherited, then whether it coalesces with the homologous genes of juvenile 1 depends on whether or not they have the same mother. If they have the same mother (which occurs with probability 1/*n*_f_), then with probability 1/2, the maternal genes of juveniles 1 and 2 are the same and all three genes coalesce with probability *F*_ST_(*t*), and with probability 1/2, the maternal genes of juveniles 1 and 2 are different and all three genes coalesce with probability *K*(*t*). If, however, juveniles have different mothers (which occurs with probability 1 − 1/*n*_f_), then for all three genes to coalesce, first the mother of juvenile 2 must be philopatric (which occurs with probability (1 − *m*_f_)) and second, three genes sampled in three different juveniles must coalesce (we call *κ*(*t*) the probability of this happening). The probability that two homologous genes of juvenile 1 and the maternally inherited gene of juvenile coalesce then reads as

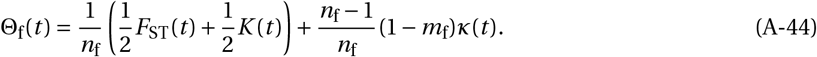

Similarly, the probability that two homologous genes of juvenile 1 and the paternally inherited gene of juvenile coalesce then reads as

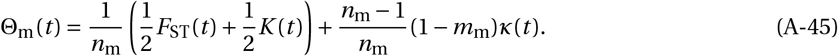

In order to compute *K*(*t* +1), we therefore need an expression for *κ*(*t*), the probability that three genes sampled in three different juveniles before dispersal coalesce. We can follow a similar argument as the one used to derive eq. (A-40). Specifically, conditioning on which genes are sampled in each juvenile (maternal or paternal), and whether these juveniles have parents in common, we obtain

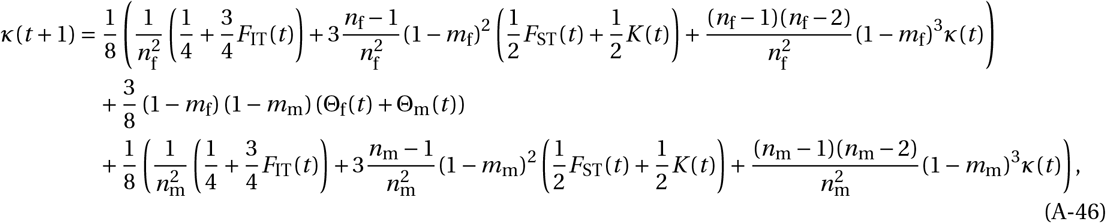

where the first line is the probability that all three genes are maternal and coalesce (decomposed according to whether the three sampled juveniles have the same mother, with probability 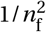, two out of three have the same mother, with probability 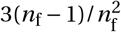, or when they have different mothers, with probability 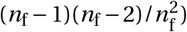; the second line is the probability that two genes are maternal and one is paternal or that one is maternal and the other two are paternal and that they coalesce; and the third line is the probability that all three genes are paternal and coalesce (decomposed according to whether the three sampled juveniles have the same father, with probability 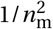, two out of three have the same father, with probability 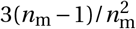, or when they have different fathers, with probability 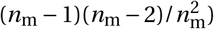.

Substituting eqs. (A-44) and (A-45) into (A-43) and (A-46) (with *F*_ST_(*t*) and *F*_IT_(*t*) at equilibrium eq. A-42) allows us to solve for the equilibrium coalescence probabilities *K = K*(*t*+1) = *K*(*t*) and *κ* = *κ*(*t*+1) = *κ*(*t*). Doing so we obtain a complicated expression for *K*, which when dispersal is weak and local population size is large is given by eq. (4) of the main text (i.e. substituting for *m*_f_ = *M*_f_/*n*_f_, *m*_m_ = *M*_m_/*n*_m_, *n*_f_ = (1 − *r*)*n* and *n*_m_ = *rn* and taking the limit *n* → ∞).

#### A.2.8 Allele frequency at a polymorphic equilibrium

In this section we derive the equilibrium frequency *p*\* of the male-detrimental, female-beneficial *A* allele when polymorphism is favoured (i.e. eq. (5) of the main text holds). By plugging main text eq. (4) (which assumes large patches) into eqs. (1)-(2) and solving for Δ*p*(*p*\*) = 0 for *p*\*, we find

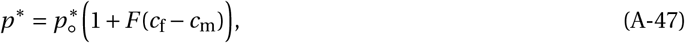

where *F* is given in eq. (4) and

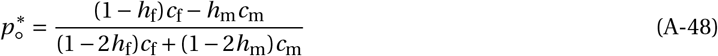

is the equilibrium frequency in a well-mixed population (see e.g. Box 3 in Mullon et al., 2012). Eq. (A-47) reveals that, compared with the well mixed case (when *F* = 0), limited dispersal (*F* > 0) leads to a greater equilibrium frequency of the female-beneficial allele when *c*_f_ > *c*_m_ (and vice-versa) and that this occurs irrespective of dominance effects (see also Figure 5A for numerical solutions of *p*\* in small patches).

### A.3 Sex-linked locus

In this section, we derive and analyse the allele frequency change at a sex-linked locus and show that the effect of limited on allele segregation is largely the same as on autosomes (our results are summarised in section 3.1.5 of the main text). Without loss of generality, we assume that our population of interest belongs to an XY species (i.e. in which males are the heterogametic sex), so that the sexually antagonistic allele segregates on the X-chromosome. All our results can be applied to ZW species by simply exchanging male and females variables.

#### A.3.1 Weighted allele frequency change

Our argument follows the same argument used for an autosomal locus in section A.2. In terms of genotype and phenotype, females are specified in the same way as for an autosomal locus (eqs. A-1 and A-3). Males however are heterogametic and therefore have genotype and phenotype respectively specified by

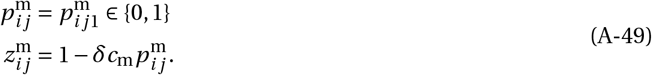

Since males only pass their X linked genes to their daughters, the weighted allele frequency change is now given by

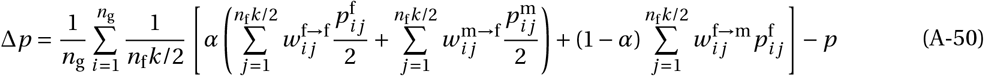

(compare with eq. A-5 for autosomes). Also due to male heterogamety, females and males now have reproductive values *α* = 2/3 and 1 − *α* = 1/3, respectively (see eq. A-4). Substituting for these reproductive values and using the fact that 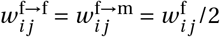 and 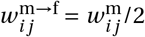 for our model, eq. (A-50) becomes

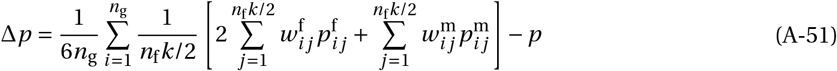

(compare with eq. A-11 for autosomes).

Substituting fitness expansions eq. (A-10) into eq. (A-52), we obtain the following expression,

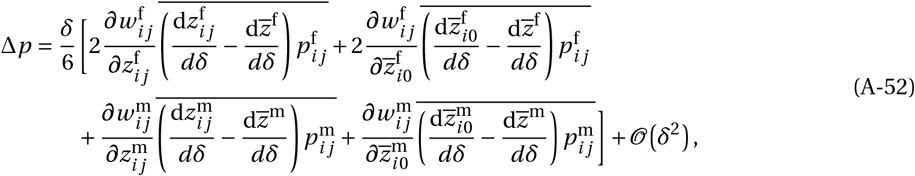

where the upper bar means the average over all juveniles and over all groups. The female related averages are computed in the same way as we did for autosomes (see section A.2.4), giving

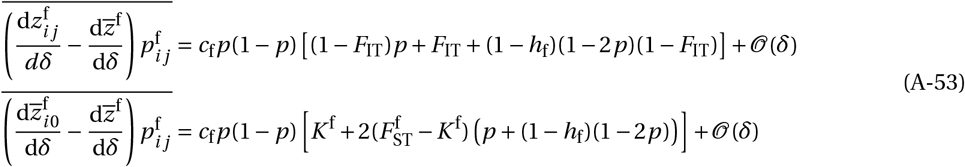

where *F*_IT_ is the probability that the two homologous gene copies at a neutral X-linked locus, sampled in a random juvenile female, are IBD (note that such a probability does not exist for males as they are heterogametic); 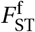 denotes the probability that two neutral X-linked are IBD when these are sampled in different female juveniles of the same group;and *K*^f^ is the probability that the two homologous gene copies in one juvenile female plus one copy in another randomly sampled juvenile female from the same group are all IBD.

For male related averages, we obtain from eq. (A-49)

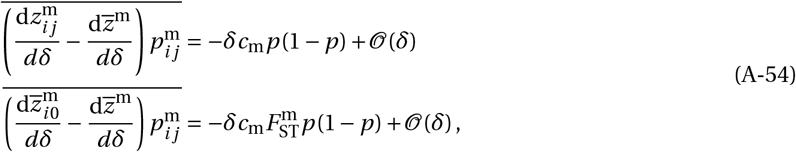

where 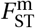 is the probability that two neutral X-linked are IBD when these are sampled in different male juveniles of the same group defined as in the preceding paragraph (note that in contrast to the autosomal case, this probability is not necessarily equal to when genes are sampled in females, i.e. 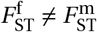, see section A.3.2 for more details).

Substituting eqs. (A-53) and (A-54) in eq. (A-52), we obtain that the change in weighted allele frequency at an X-linked locus is

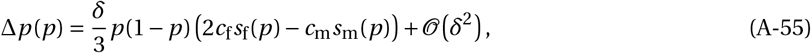

where

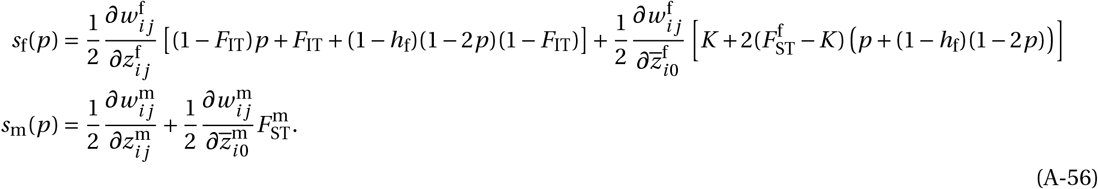

In turn, plugging fitness derivatives eq. (A-31) into eq. (A-56), we obtain

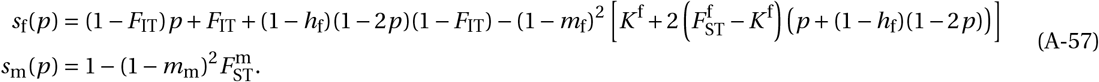

While the general form *s*_f_(*p*) is unchanged from the autosomal case (see eq. 2 of main text), *s*_m_(*p*) no longer includes direct or indirect effects arising from male heterozygotes as male hemizygosity precludes the existence of male heterozygotes. Thus, the effect of selection in males on allele frequency change, *c*_m_ × *s*_m_(*p*) in eq. (A-55), depends only on *c*_m_ and the strength of male kin competition (determined by 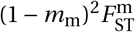 in eq. A-57). To analyse allele dynamics at an X-linked locus further, we compute the relevant coalescence probabilities in the next section.

#### A.3.2 Coalescence probabilities at a neutral X-linked locus

In this section, we calculate the coalescence probabilities that are relevant for selection (eq. A-56) at an X-linked locus, using similar arguments as those used for an autosomal locus (see section A.2.7).

##### Pairwise coalescence probabilities

Let us first consider the probabilities that two neutral X-linked genes sampled at generation *t* + 1 in two neighbouring juvenile females, 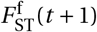, and males, 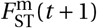, coalesce. As for the autosomal case (see eq. A-36), we can decompose 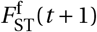 according to whether the two sampled genes in females are either both of maternal or paternal origin (in which case they coalesce with respective probabilities *θ*_f_(*t*) and *θ*_m_(*t*)) or one is of maternal and the other of paternal origin (in which case they coalesce with respective probabilities *θ*_f,m_(*t*) and *θ*_m_(*t*)). For X-linked genes sampled in two different males, 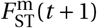, both genes are necessarily of maternal origin. We thus obtain a recurrence of the form

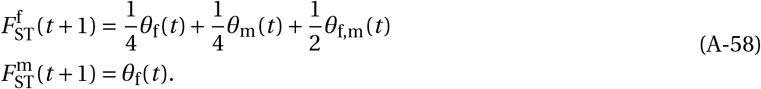

Then, coalescence probabilities according to parental origin can be decomposed as

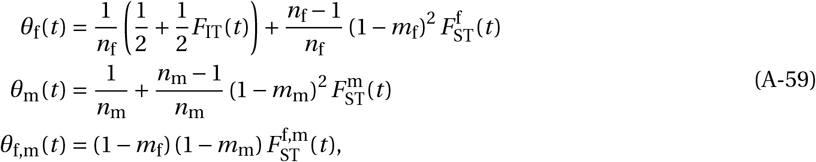

where *F*_IT_(*t*) is the probability that the two homologous gene copies at a neutral X-linked locus in a randomly juvenile female at generation *t* coalesce, and 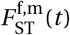 is the coalescence probability for two X-linked genes, one sampled in a juvenile male and another in a juvenile female (from the same group) at generation *t*. In turn, these two probabilities satisfy,

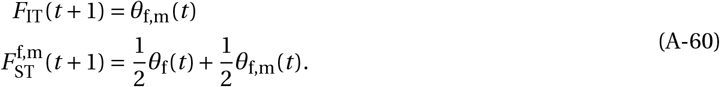

Plugging eq. (A-59) into eqs. (A-58) and (A-60) and solving simultaneously for the equilibrium 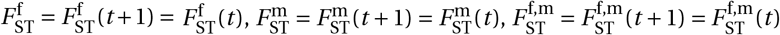, and *F*_IT_ = *F*_IT_(*t* + 1) = *F*_IT_(*t*) allows us to specify the relevant coalescence probabilities for selection on an additive allele (*h*_f_ = 1/2) in terms of demographic parameters and sex-specific dispersal. Because such a calculation leads to complicated (but analytical) expressions, we only present their approximation when patches are large and dispersal is weak in the main text (by substituting for *m*_f_ = *M*_f_/*n*_f_, *m*_m_ = *M*_m_/*n*_m_, *n*_f_ = (1 − *r*)*n*, and *n*_m_ = *rn* and taking the limit *n* → ∞,in which case we find 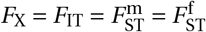 given by eq. (7) in the main text, which is equivalent to eq. 9 of Ramachandran et al., 2008).

##### Threeway coalescence probabilities

We now develop expressions for threeway coalescence probabilities. Recall that for selection (eq. A-56), we seek to compute *K*^f^, which is the probability that the two homologous gene copies in one juvenile female plus one copy in another randomly sampled juvenile female from the same group are all IBD. We further introduce *K*^m^, which is the probability that the two homologous gene copies in one juvenile female plus one copy in a randomly sampled juvenile male from the same group are IBD, and which will be necessary in our calculations. Using a similar argument as for the autosomal case (see eq. A-43), we can write,

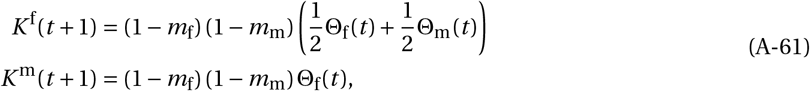

where Θ_f_(*t*) is the probability that all three relevant genes coalesce when the one gene sampled as a single copy is maternally inherited, and Θ_m_(*t*) when it is paternally inherited. These probabilities can be expressed as

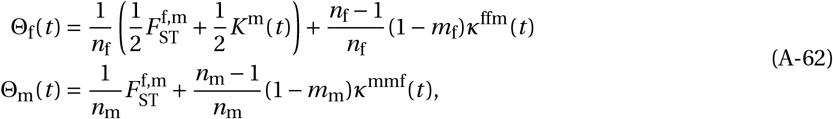

where *κ*^ffm^(*t*) is the probability that three genes, randomly sampled in two different female and one male juvenile from the same group all coalesce, and *κ*^mmf^(*t*) is the coalescence probability for genes randomly sampled in two different male and one female juvenile from the same group. In turn, such probabilities can be written as

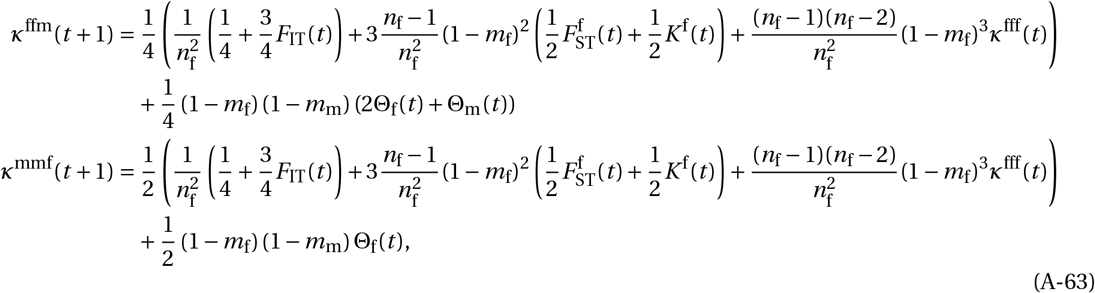

which depend on *κ*^fff^(*t*) and *κ*^mmm^(*t*), the coalescence probabilities for genes randomly sampled in three different female and male juveniles, respectively. In turn, these two latter probabilities themselves satisfy the recurrences

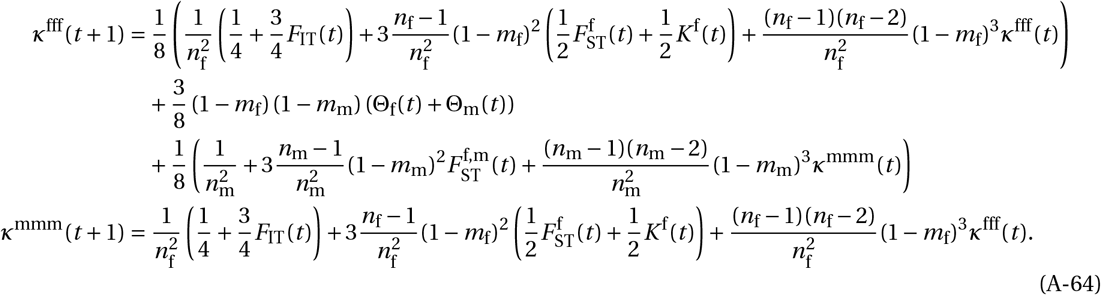

Substituting eqs. (A-62) into eqs. (A-61), (A-63) and (A-64) finally yields a dynamical system whose equilibrium we can solve to obtain the relevant quantity, *K*^f^ = *K*^f^(*t* + 1) = *K*^f^(*t*), for selection. In the high density-low migration limit (letting *m*_f_ = *M*_f_/*n*_f_, *m*_m_ = *M*_m_/*n*_m_, *n*_f_ = (1 − *r*)*n* and *n*_m_ = *rn* and taking the limit *n* → ∞), we obtain

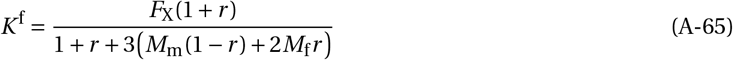

where *F*_X_ is given by eq. (7) in the main text.

#### A.3.3 Analysis of allele frequency change at a sex-linked locus

In this section, we analyse eqs. (A-55)-(A-57) using eqs. (7) and (A-65) assuming first additive and then nonadditive allele effects, in order to understand the effects of limited dispersal on the segregation of sexually antagonistic alleles on the X-chromosome.

##### Additive effects

To obtain eq. (6) of the main text, we substitute eq. (7) into eqs. (A-55)-(A-57) with *h*_f_ = 1/2 and *c*_f_ = *c*_m_ = *c*.

##### Non-additive effects

Plugging eqs. (7) and (A-65) into eqs. (A-55)-(A-57) and deriving conditions such that Δ*p*′(0) > 0 and Δ*p*′(1) < 0, we find that the conditions favouring the maintenance of polymorphism under non-additive effects are

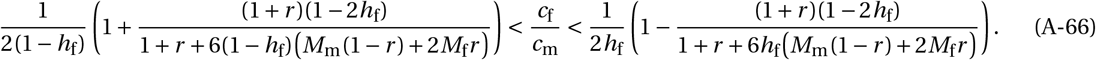

Note that if males and females are not dispersal limited (i.e. when *M*_m_ and *M*_f_ tend to infinity), eq. (A-66) reduces to the polymorphism condition of a well-mixed population (e.g. eq. 2 in Fry, 2010, under weak selection). However, as increasingly limited dispersal leads to inbreeding and a dearth of female heterozygotes, the polymorphism space contracts (see left and right sides of the inequality A-66 as *M*_m_ and *M*_f_ decrease). Furthermore, because coalescence occurs faster on the X-chromosome than autosomes (main text eqs. 4 and 7), this narrowing of polymorphism conditions occurs more rapidly for X-linked sexually antagonistic loci (compare Figure 3A and B Top).

As with autosomes, sex-biased dispersal at X-linked loci leads one sex to experience stronger kin competition and shifts the polymorphism space in favour of stronger antagonistic effects in the less dispersive sex (Figure 3B Bottom). Notice, however, that the size of polymorphism conditions on the X-chromosome are affected by the immigrant sex ratio even when there are equal numbers of males and females (*r* = 0.5), becoming smaller when immigrants are male-biased. This is because coalescence probabilities (and thus frequency of inbreeding) on the X-chromosome are increased when males are the more dispersive sex (eq. 7).

Finally, solving for Δ*p*(*p*\*) = 0 (from eqs. A-55-A-57 with eqs. 7 and A-65), we find that when eq. (A-66) holds and selection favours polymorphism, the equilibrium allele frequency *p*\* on an X-chromosome is given by

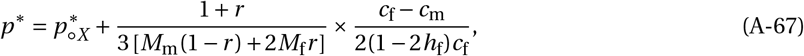

where

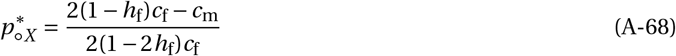

is the equilibrium in a well mixed population. Inspection of the last term of eq. (A-67) reveals that *p*\* will increase or decrease depending on whether the sign of *C*f-*c*_m_ is positive or negative respectively. Thus, eq. (A-67) shows that, relative to a well-mixed population, limited dispersal (low values of *M*_m_ and *M*_f_) leads selection to favour the sex which feels the greatest cost of their detrimental allele independently of dominance (in this case *h*f), like on autosomes (see eq. A-47).

## B Individual based simulations

We simulated a diploid population subdivided among *n*_g_ = 50 groups following the life-cycle elaborated in Appendix A.1 (our simulation code can be downloaded from the). Specifically, at the beginning of a generation, adult females in each patch produce large poisson distributed number (mean *k* = 20) of eggs. Such high female fecundity effectively precludes the possibility of patch extinction from stochastic fluctuations in the number of eggs being laid. Each egg is fertilised by a randomly sampled male within the same patch to produce a zygote whose sex is determined at random (with probability 1/2 to be male and 1/2 to be a female). Each zygote disperses independently with sex-specific dispersal probability *m*_m_ and *M*_f_ for males and females, respectively. If a zygote disperses, it joins a patch other than its natal one at random (so each remaining patch has a probability 1/(*n*_g_ − 1) of being the destination of dispersal). After dispersal, zygotes within each patch compete intrasexually to form the new adult mating pool. To model this, *n*_m_ males and *n*_f_ females are sampled without replacement from the male and female zygotic pools in each patch, where the probability of an individual being sampled is weighted by the competitiveness of its genotype (according to Figure 1A).

Simulations were initiated with both alleles at frequency 0.5 at Hardy-Weinberg equilibrium across the whole population and ran until one allele was fixed. 10,000 replicates were conducted for each parameter combination. For Figure 4, we varied proportion of immigrants: (*M*_m_ + *M*_f_)/(*n*_m_ + *n*_f_) = 10^-2.7^,10^-2.6^,…,10^-0.1^,1, the strength of selection: *c = c*_m_ = *c*_f_ = 0,0.05,0.2 and dominance effects: *h*_m_ = *h*_f_ = 0.2,0.5. Meanwhile for Figure 5 we varied the immigrant ratio: *M*_m_/(*M*_m_ + *M*_f_) = 0.1,0.3,0.5,0.7,0.9, and the ratio of homozygous allele costs: *c*_f_/*c*_m_ = 0.25,0.82,1,1.22,4 (for other simulation parameters, see figure legends). At the end of each generation, we calculated *F*_IT_ as

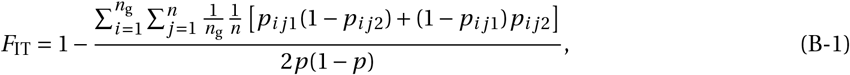

where *n* = *n*_m_ + *n*_f_ is the number of individuals of either sex in a given group, *p*_*ij*1_ and *p*_*ij*2_ is the frequency of *A* in individual indexed *j* of group *i* at the maternally and paternally inherited locus respectively, and 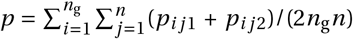 is the frequency of allele *A* in the whole population (so the fraction in eq. B-1 is the ratio of the observed number of heterozygotes to the expected number of heterozygotes under Hardy-Weinberg equilibrium). *F*_ST_ is calculated as the ratio of the genetic covariance between individuals within groups to half the expected total genetic variance in the population at Hardy-Weinberg equilibrium, i.e. as

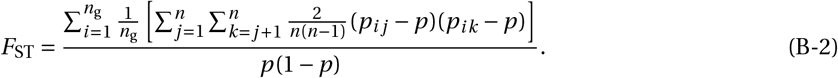

Eqs. B-1 and B-2 follow from eqs. A-20 and A-21 respectively, but are calculated in adults. To generate the points in Figure 4B, we averaged *F*_IT_ and *F*_ST_ calculations across all generations in a simulation run, with the exception of the last five (as one allele would be very close to fixation), and then across all replicate runs of a given parameter combination.

**Supplementary Figure 1:**
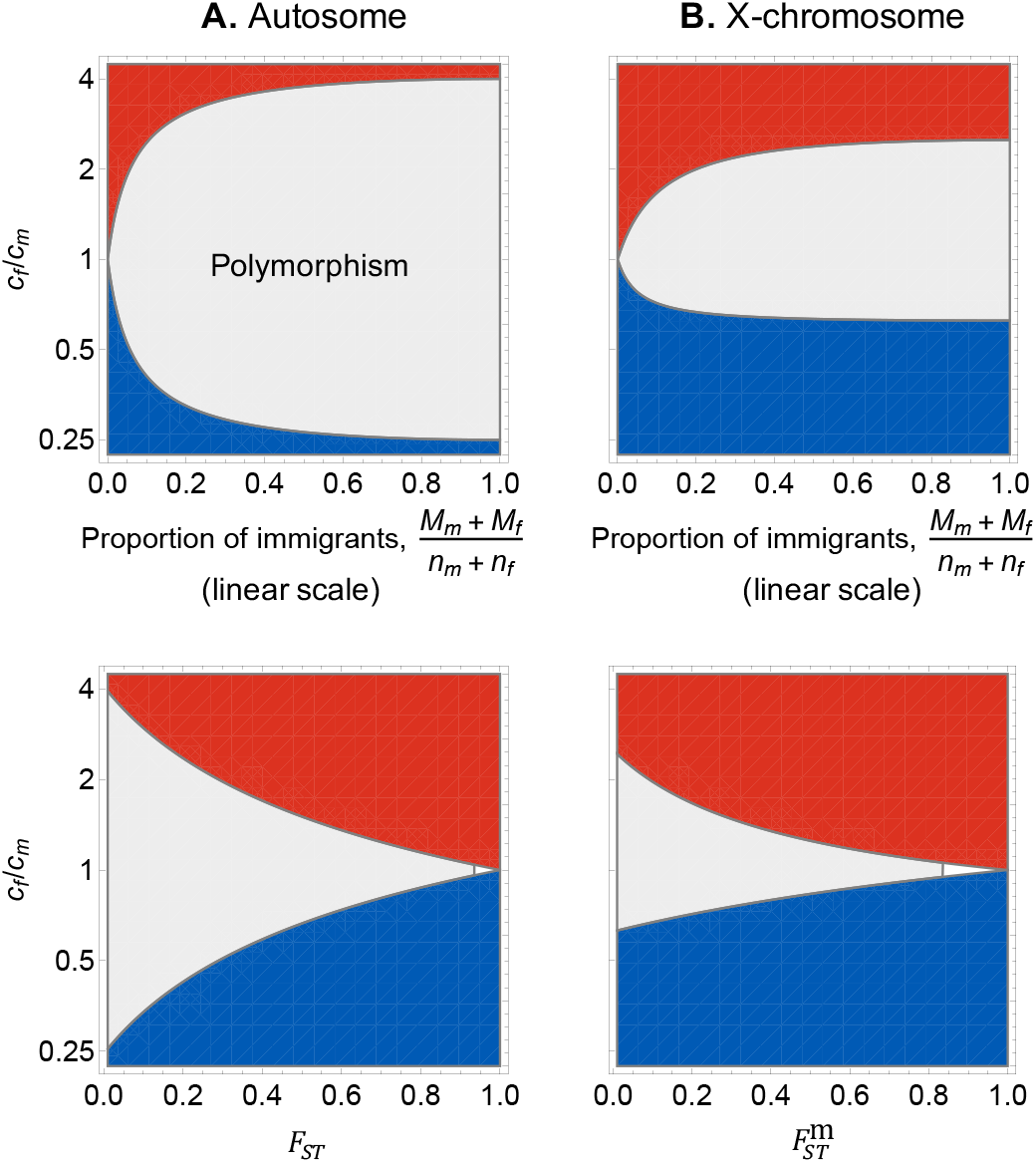
Parameters that favour the maintenance of sexually antagonistic variation and association with *F*-statistics. Combinations of parameters that lead to either balancing selection (grey, when Δ*p*′(0) > 0 and Δ*p*′(1) < 0), positive selection for *A* (red, when Δ*p*′(*p*) > 0 for all *p*), and positive selection for *a* (blue, when Δ*p*′(*p*) < 0 for all *p*) at **A.** an autosomal locus (computed from eqs. 1-2 with exact coalescence probabilities, see Appendix A.2.7 for calculations of these probabilities) and **B.** X-linked locus (computed from eqs. A-55-A-57 with exact coalescence probabilities, see Appendix A.3.2 for calculations). Top: selection according to ratio of homozygotic effects in females and males (*c*_m_/*c*_f_) and the expected proportion of immigrants in each patch at each generation ((*M*_m_ + *M*_f_)/(*n*_m_ + *n*_f_) (on a linear scale);other parameters: *n*_m_ = *n*_f_ = 5, *h*_f_ = *h*_m_ = 0.2, *M*_m_ = *M*_f_). Bottom: selection according to ratio of homozygotic effects in females and males (*c*_m_/*c*_f_) and relevant *F*_ST_ (here shown among males for the X-chromosome) when patches are large (*n*_m_ + *n*_f_ = 100) and dispersal is symmetric among the sexes (*M*_m_ = *M*_f_; other parameters: *h*_f_ = *h*_m_ = 0.2). As described in section 3.1.3, the parameter conditions which favour polymorphism become more restrictive with limited dispersal, and therefore with increasing *F*_ST_, on both autosomes and X-chromosomes. Note that this effect also occurs regardless of whether group size is small or large (top versus bottom panels), as the most relevant quantities for determining genetic structure are the effective numbers of male and female immigrants in a patch (*M*_m_ and *M*_f_, see eqs. 4 and A-42).

**Supplementary Figure 2:**
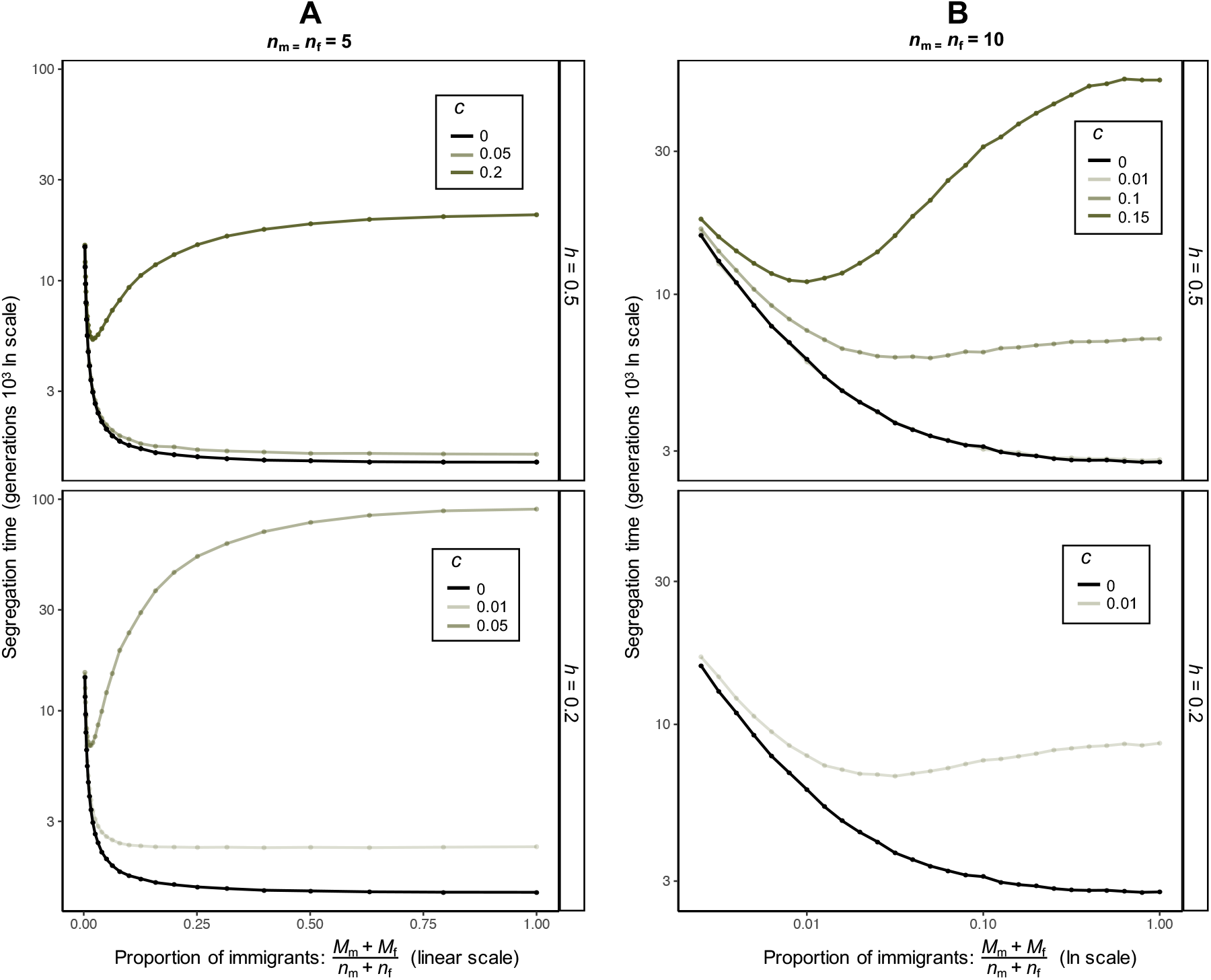
Additional plots showing allele segregation time in individual-based simulations. Segregation time averaged across replicate runs as a function of total fraction of immigrants, selection strength (*c* = *c*_f_ = *c*_m_), and dominance regime (*h = h*_f_ = *h*_m_). **A.** Shows segregation time for small groups (*n*_m_ = *n*_f_ = 5, as in Figure 4) on a linear scale (black line shows neutrality, green lines show, from lightest to darkest, *c* = 0.01, *c* = 0.05 and *c* = 0.2). **B.** Shows segregation time for larger groups (*n*_m_ = *n*_f_ = 10) on a natural log scale (black line shows neutrality, green lines show, from lightest to darkest, *c* = 0.01, *c* = 0.1 and *c* = 0.15, note that black and lightest green lines overlay each under when *h* = 0.5). See Appendix B for more details on simulation procedure.

